# Evolution of transposons as controlling elements

**DOI:** 10.64898/2026.08.24.746577

**Authors:** Taku Sasaki, Carles Borredá, Sota Fujii, Seiji Takayama, Leandro Quadrana

## Abstract

Transposons were originally described as “controlling elements,” yet the molecular basis and evolutionary origin of their transcriptional regulatory activities remain elusive. Here, we show that transcriptional regulatory activity is an intrinsic property of transposases (TPases). We demonstrate that the Arabidopsis *AtMu1* TPase is sufficient to induce sequence-specific transcriptional activation of cognate copies, revealing a pre-existing *cis*-regulatory network. We further demonstrate that long-recognized *Spm* transposon-encoded transcriptional regulator TnpA evolved from an ancestral TPase, linking transcriptional regulators to TPases. Across eukaryotes, we identify multiple domesticated TPase-derived proteins that recurrently lack catalytic residues for transposition, consistent with a one-step regulatory co-option through loss of catalytic activity. Together, our results reveal TPases as autonomous sequence-specific transcriptional regulators and provide a direct evolutionary route from TPases to transcription factors.

## Main Text

Transposons are ubiquitous genetic elements capable of mobilizing themselves through self-encoded transposases (TPases) that recognize element-specific terminal sequences within cognate transposons and catalyze excision and reintegration. Although transposition threatens genome integrity, the long-term conflict between transposons and host defense systems has significantly contributed to the evolution of genomes and cellular pathways (*1*, *2*). Transposons have also emerged as major drivers of transcriptional innovation because they frequently carry transcription factor (TF)-binding sites that, upon transposition, redistribute *cis*-regulatory elements throughout the genome and rewire transcriptional networks (*3*-*7*). Such transposon-mediated rewiring has played central roles in the evolution of diverse biological processes, including zygotic genome activation and the interferon response (*8*, *9*). Beyond dispersing TF-binding sites, transposons have themselves contributed to the evolution of TFs. TPases possess sequence-specific DNA-binding activity, a defining property of TFs, and have repeatedly been domesticated into host transcriptional regulators during evolution. In many cases, co-option involved fusion of a TPase-derived DNA-binding domain (DBD) with a host regulatory domain (*10*-*12*). Yet some TFs appear to derive directly from TPases, leaving unresolved how transcriptional regulatory activity arose in the absence of such a fusion.

Barbara McClintock conceptualized transposons as “controlling elements” based on their ability to influence gene expression. Classical genetic studies in maize showed that diverse transposons, including *Ac* (*Activator*), *Spm* (*Suppressor-Mutator*), and *Mu* (*Mutator*), modulate the expression of nearby genes in a transposon activity-dependent manner (*13*-*18*). However, these pioneering studies, largely based on genetic analysis at individual loci, did not reveal the molecular basis and genome-wide principle underlying this regulatory activity. Recent studies have identified transposon-encoded factors that mediate sequence-specific transcriptional activation through epigenetic reprogramming. In Arabidopsis, *Mu*-like *VANDAL* transposons encode the accessory protein VANC, which functions as a sequence-specific transcriptional regulator that activates cognate *VANDAL* sequences (*19*-*21*). Genome-wide analyses further predicted additional transposon families with similar epigenetic regulatory activities, including Arabidopsis *Spm*, suggesting that transposon-encoded transcriptional regulators may be widespread (*21*). The maize *Spm* transposon encodes TPase (TnpD) and an accessory regulator (TnpA) that are both required for transposition (*22*-*25*). In addition, TnpA has been linked to epigenetic and transcriptional changes (*26*, *27*), although its origin, genome-wide impact, and mechanisms of action are unknown.

### Sequence-specific TnpA regulatory systems in Arabidopsis

To identify transposon-mediated transcriptional regulatory systems, we focused on Arabidopsis *Spm* elements, whose accessory factor TnpA has been implicated in transcriptional and epigenetic regulation in maize and Arabidopsis (*21*, *26*, *27*), and harbors mobile copies in Arabidopsis (*28–30*). To this end, we developed an EGFP-based reporter system based on the *Spm1* element AT2TE22040, which shows the strongest transcriptional activity in DNA hypomethylated mutants (*31*). The internal sequence of AT2TE22040 was replaced with EGFP and transformed into wild-type (WT) plants (Spm1-EGFP, fig. S1A). Although Spm1-EGFP was epigenetically silenced in WT background, it was *trans*-activated when crossed with *ddm1* mutant plants, which show broad transposon activation, indicating the presence of a *trans*-acting regulatory factor for *Spm1* (fig. S1, supplementary text 1). By linkage mapping analysis, we identified the endogenous *Spm1* copy responsible for Spm1-EGFP reporter activation (supplementary text 1). We designated this copy “*KAKUSEI*” (“awakening” in Japanese). Transgenic expression of the full-length *KAKUSEI* restored EGFP expression from the epigenetically silenced Spm1-EGFP reporter (Fig. 1A and B). *KAKUSEI* harbors a single gene encoding a factor related to maize TnpA (fig. S1H), hereafter referred to as TnpA^Spm1^.

**Fig. 1.**
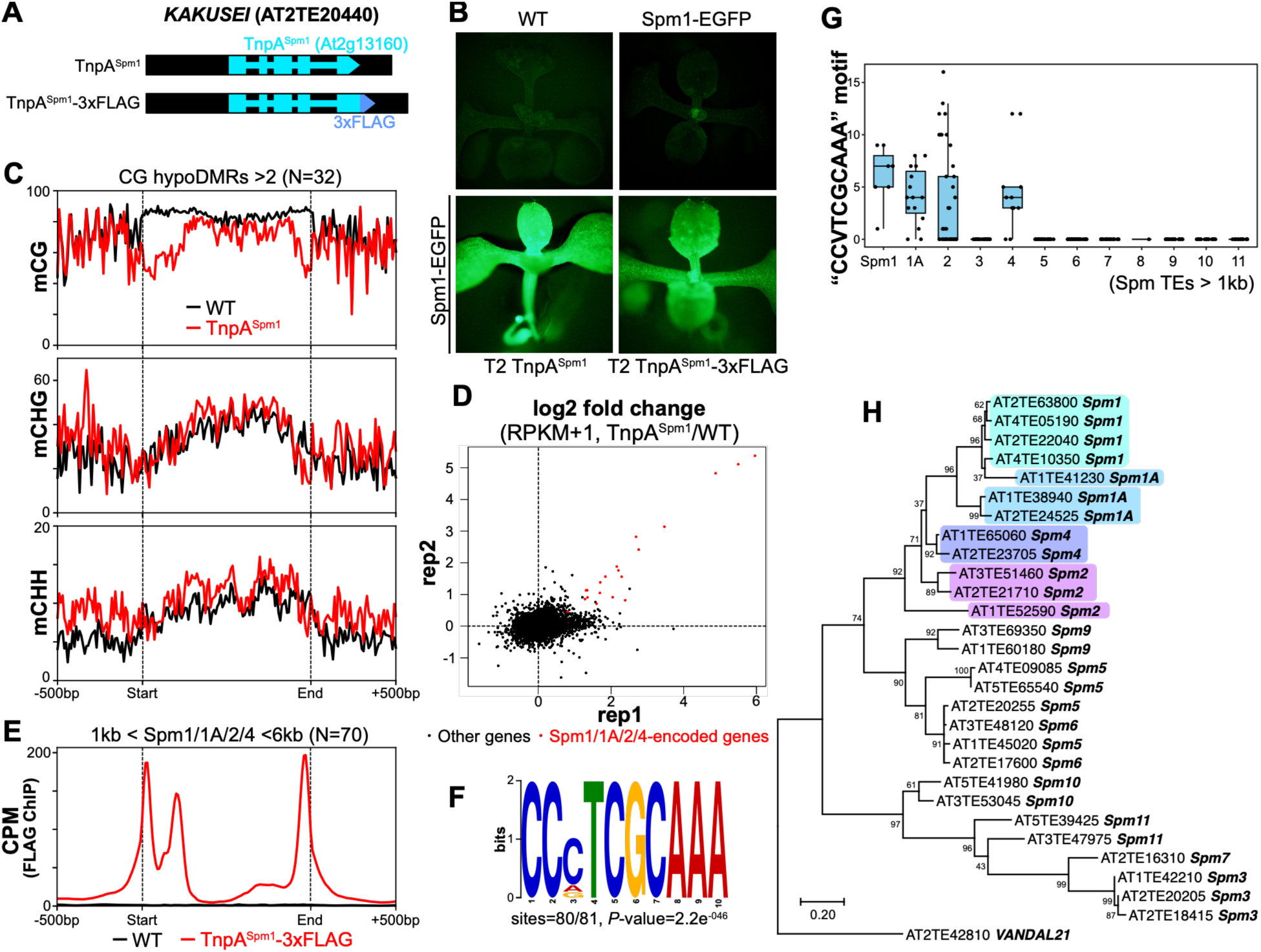
Characterization of novel transposon-encoded transcriptional regulator in *Spm* transposons. (**A**) Schematic representation of AT2TE22040 (*Spm1*, *KAKUSEI*) and its FLAG-tagged version. Blue and black regions indicate coding and noncoding regions within the transposon, respectively. (**B**) EGFP fluorescence in seedlings observed under a fluorescence microscope. (**C**) Metaplot of DNA methylation levels across *Spm1*, *1A*, *2*, and *4* transposons (1-6 kb in length, N=70), including 500 bp flanking regions. DNA methylation in WT and TnpA^Spm1^ is shown in black and red lines, respectively. Vertical dashed lines indicate transposon boundaries. (**D**) Scatter plot of RNA-seq results from two replicates. Each dot represents the log2 fold change in transcript level (RPKM+1, TnpA^Spm1^/WT). Genes encoded within *Spm1*, *1A*, *2*, and *4* are shown in red and others shown in black. (**E**) Metaplot of ChIP-seq signals (CPM; counts per million reads) for 3xFLAG-TnpA^Spm1^ across *Spm* transposons containing ≥2 CG hypoDMRs (N=32). Black and red lines represent WT and 3xFLAG-TnpA^Spm1^, respectively. (**F**) Putative target motif of TnpA^Spm1^ identified from ChIP-seq peaks within *Spm* transposons (N=80) using DREME of MEME suite. (**G**) Boxplot showing the number of “CCVTCGCAAA” (V=A, C, or G) motifs within *Spm* transposons (>1 kb in length). Each dot represents a single transposon. (**H**) Phylogenetic tree of *Spm* transposons in Arabidopsis, based on sequence alignment of the terminal 120 bp from 5’ ends.

To assess the global effect of TnpA^Spm1^ on DNA methylation, we performed methylome analysis of plants expressing TnpA^Spm1^ as well as control WT plants. Both termini of endogenous *Spm1* transposons show strong DNA hypomethylation in TnpA^Spm1^ plants compared to WT, with CG methylation being most strongly affected (Fig. 1C, fig. S2). Unlike *VANDAL* transposons, where anti-silencing factor VANCs cause spreading of non-CG hypomethylation (*19*-*21*, *32*), TnpA^Spm1^-induced DNA hypomethylation is restricted to transposon termini, with internal DNA methylation largely unchanged (Fig. 1C, fig. S2C and D). Different degrees of hypomethylation were also observed in related *Spm* families, including *Spm1A*, *2*, and *4* (fig. S2B to E). Notably, heterochromatic histone modifications such as H3K9me remained at target *Spm* elements despite their activation by TnpA^Spm1^ (fig. S3A and B). We further examined the underlying pathway involved in TnpA^Spm1^-mediated hypomethylation and found that host-encoded active DNA demethylation catalyzed by the DNA glycosylases ROS1, DML2, and DML3 is required (fig. S3, supplementary text 2). These results indicate that TnpA^Spm1^-mediated hypomethylation hijacks host DNA demethylation pathways. RNA-seq analysis revealed that TnpA^Spm1^-induced hypomethylation was accompanied by transcriptional activation of genes encoded within these transposons (Fig. 1D), consistent with the observed induction of Spm1-EGFP reporter. We next used chromatin immunoprecipitation followed by sequencing (ChIP-seq) to determine TnpA^Spm1^ target recognition. We generated transgenic plants expressing FLAG-tagged TnpA^Spm1^ (TnpA^Spm1^-3xFLAG), which we confirmed to retain transcriptional regulatory function (Fig. 1B). ChIP-seq analysis revealed that TnpA^Spm1^ binds the termini of *Spm1*, *1A*, *2*, and *4*, coinciding with TnpA^Spm1^-induced hypomethylated regions (Fig. 1C and E, fig. S4A to C). TnpA^Spm1^-binding regions were strongly enriched with the “CCVTCGCAAA” motifs (V: A, C, or G) (Fig. 1F), which were concentrated in these related four families forming a monophyletic group (Fig. 1G and H). Although the Arabidopsis reference genome (TAIR10) contains 729 such motifs, regions outside *Spm* transposons generally show low motif density (≦2 motifs/kb) and little or no TnpA^Spm1^ binding (fig. S4D). A notable exception was a non-*Spm* pericentromeric high-density motif cluster bound by TnpA^Spm1^ (fig. S4E to F, supplementary text 3). Altogether, these results indicate that high-density motif clusters are essential for target-specific transcriptional regulation by TnpA^Spm1^.

Analysis of DNA methylation in epigenetic recombinant inbred lines (epiRILs), which segregate chromosomal regions with distinct epigenetic states (*33*, *34*), revealed strong *trans* DNA hypomethylation activities for the 12 *Spm* families annotated in the Arabidopsis genome and identified family-specific sequence motifs associated with *trans* DNA hypomethylation targeting, suggesting co-diversification of target specificity and transposon lineages (fig. S5, supplementary text 4). Further experimental characterization of the mobile *Spm3* family (*28*-*30*) identified a TnpA^Spm3^ factor that induces sequence-specific hypomethylation at cognate copies, demonstrating that multiple TnpA-mediated regulatory systems exist across diverse *Spm* transposons (fig. S5, supplementary text 4).

Together, these findings establish TnpA as a sequence-specific regulator that binds clustered cognate motifs and directs locus-specific epigenetic and transcriptional activation. Like canonical TFs, TnpA establishes a *cis*-regulatory network by targeting dispersed *Spm* copies across the genome, providing a molecular basis for transposon-encoded transcriptional regulation.

### A transposon-encoded regulator rewires host gene expression

Transposons or their derivatives can influence the transcription of nearby genes (*35*-*38*). We next investigated whether TnpA^Spm1^ functions as a controlling element for host genes. To directly test this possibility, we developed a synthetic reporter based on the *SUPPRESSOR OF DRM1 DRM2 CMT3* (*SDC*) locus (Fig. 2A). In WT plants, *SDC* is transcriptionally silenced by DNA methylation of tandem repeats located within its promoter. Loss of promoter methylation, as observed in the *ddc* (*drm1 drm2 cmt3*) mutant, activates *SDC* expression and produces a characteristic curled-leaf phenotype (*36*) (Fig. 2B). We generated a dSpm1-SDC construct by replacing these repeats with an engineered non-coding *Spm1* fragment containing the TnpA^Spm1^-targeted motifs. Transgenic dSpm1-SDC plants developed normally, consistent with silencing of *dSpm1* sequence and *SDC* repression. By contrast, induction of TnpA^Spm1^ in dSpm1-SDC plants triggered strong overexpression of *SDC* accompanied by the curled-leaf phenotype (Fig. 2B and C). Together, these results establish Arabidopsis transposon-encoded transcriptional regulators as mechanistic drivers of host gene regulation through cognate transposon sequences, providing a molecular basis for McClintock’s “controlling element” concept.

**Fig. 2.**
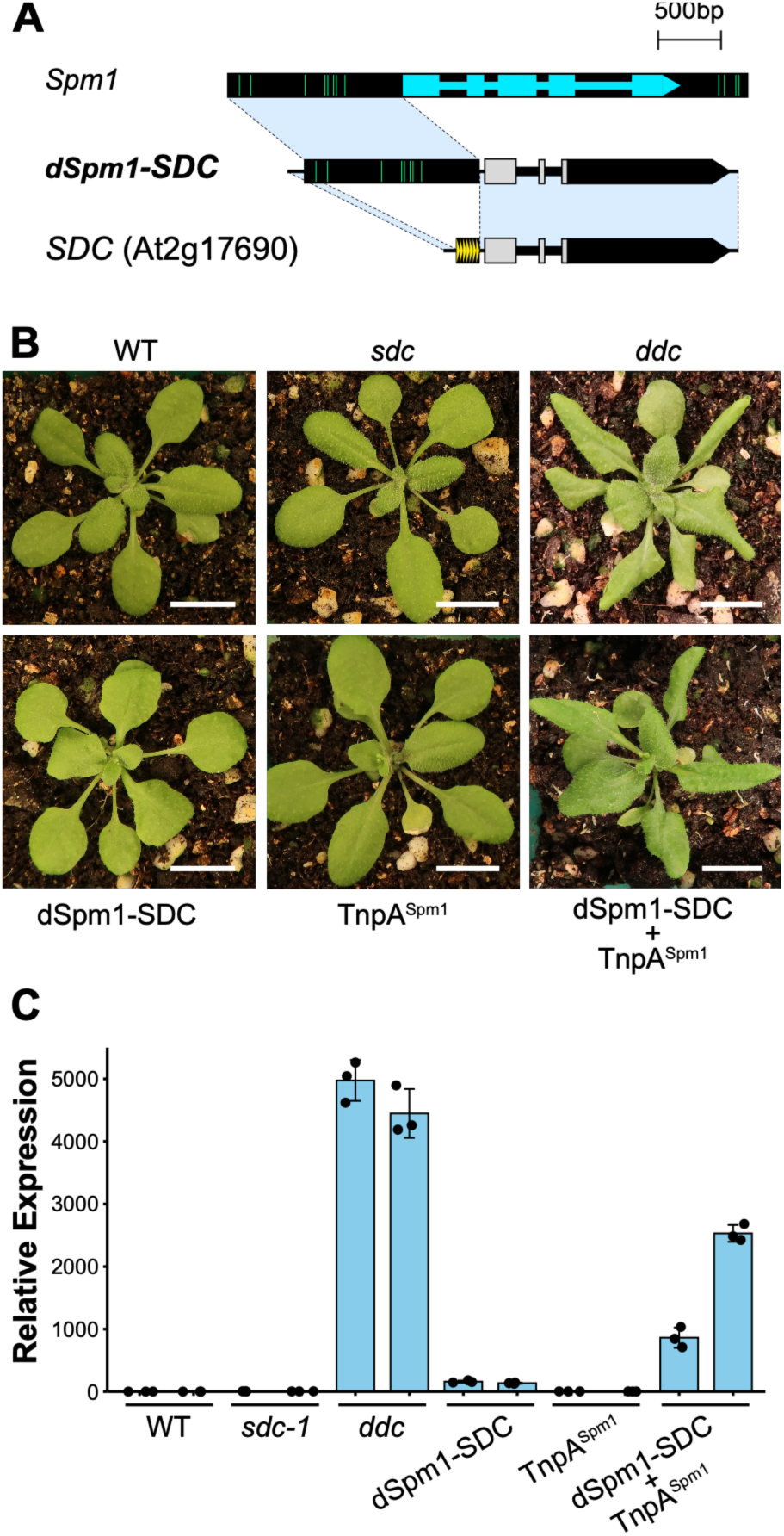
Transcriptional regulatory activity of TnpA^Spm1^. (**A**) Schematic representation of *SDC* gene (At2g17690) and the dSpm1-SDC reporter construct. Tandem repeats in the *SDC* promoter were replaced with a defective Spm1 (dSpm1) element consisting of the 5’ noncoding region of *KAKUSEI*. (**B**) Leaf morphology of seedlings for each genotype. White horizontal bars indicate 1 cm. The DNA hypomethylated *ddc* mutant was used as a positive control for *SDC* overexpression. (**C**) Relative transcript levels of *SDC* gene determined by RT-qPCR. Expression levels were normalized to those of *UBC* gene and are shown relative to WT. Error bars represent the standard deviation from three technical replicates.

### Evolution of Spm-encoded regulators from TPases

To explore the evolutionary origin of this transposon-encoded regulatory activity, we performed structure-based comparisons (*39*) of 962 predicted protein structures encoded by reference *Spm* transposons from a wide range of eukaryotes, including plants, animals, and fungi. As expected, the core catalytic domain of TPases showed structural conservation across diverse taxa (Fig. 3A). In addition, we identified various accessory proteins and domains unrelated to the catalytic TPase core, including numerous TnpA-like proteins, which appear to be restricted to flowering plants (Fig. 3A). Further analyses indicated that TnpA-like proteins, including TnpA^Spm1^ and TnpA^Spm3^ (*40*), contain a DBD composed of two tandem tri-helical helix-turn-helix (HTH) motifs separated by about 45 amino acids (Fig. 3B). This HTH architecture is also present in other accessory proteins and TPases from distantly related eukaryotes (Fig. 3A to C), suggesting that the TnpA DBD originated from an ancestral TPase. Consistent with this model, the DBD is typically fused to the TPase core in fungi and animals but is encoded separately, either in alternative isoforms or as separated genes, in plant *Spm* elements (Fig. 3C). Analyses of protein domain organization and exon-intron structure across 78 *Spm* elements confirmed that DBDs are commonly flanked by disordered linker peptides and exon-exon splice junctions (Fig. 3D). To further investigate this organization, we analyzed long-read transcriptomic data from zebrafish embryos and Arabidopsis *ddm1* mutant, conditions in which transposons become transcriptionally active (*41*, *42*). These data confirmed the partitioning of TPases and DBDs across distinct exons, further supporting the emergence of TnpA-like proteins through alternative splicing of TPases (Fig. 3E). In addition, we identified numerous TnpA-like proteins encoded by separate genes in Arabidopsis (Fig. 3E). Cap Analysis of Gene Expression (CAGE) mapping of transcriptional start sites confirmed these are transcribed independently (fig. S6A), indicating that TnpA-like factors and TPases have evolved to be encoded by separate genes. Phylogenetic analyses of *Spm* further support that ancestral TPases contained the DBD (fig. S6B). Together, these findings support the evolution of *Spm*-encoded transcriptional regulatory factors from ancestral TPases through alternative splicing and functional domain partitioning.

**Fig. 3.**
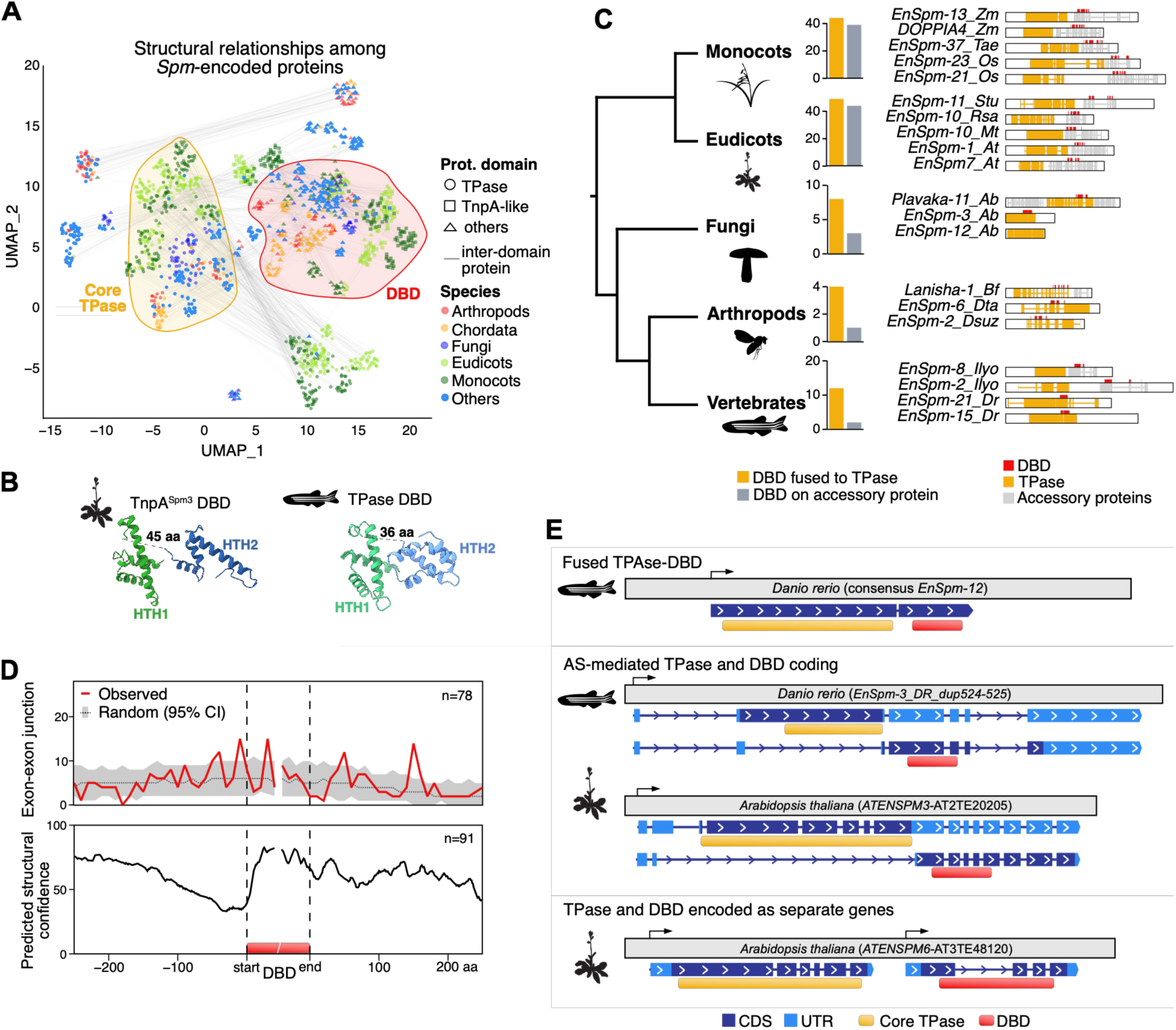
Evolution of transcriptional regulators through functional separation from TPases. (**A**) UMAP representation of the structural domains encoded by MuDR proteins. Domains belonging to the same protein are connected by lines. Each domain was classified as a TPase, or TnpA-like, or others based on structural alignments. (**B**) Structural comparison of the DBDs from the Arabidopsis TnpA^Spm3^ (AT2TE20205) and *Danio rerio* EnSpm-21 consensus TPase protein. Linker peptides between both HTH motifs are represented by dashed lines. (**C**) Localization of the DBDs relative to the TPase across major eukaryotic taxa. Representative examples for each taxon are shown on the right. (**D**) Metaplot of exon-exon junctions and pLDDT scores centered on the DBD. For exon-exon junctions, the expected random distribution and its 95% confidence interval are also displayed. (**E**) Representative examples of distinct DBD-TPase architectures. Top: DBD and TPase are fused into a single protein. Middle: Alternative splicing of a single gene generates transcript encoding either the DBD or the TPase. Bottom: DBD and TPase are encoded by separate genes. Transcription start sites are represented by arrows.

### Transcriptional regulatory ability is intrinsic to TPase

The origin of the TnpA DBD from an ancestral TPase raised the possibility that transcriptional regulatory activity preceded its functional separation. To test whether regulatory activity can be encoded by a TPase itself, we sought a transposition-competent transposon lacking accessory factors. Among the DNA transposons that retain transposition activity in Arabidopsis (*28*, *43*, *44*), *VANDAL21* and *Spm3* encode accessory regulators (*19*-*21*, this study), whereas *AtMu1* contains only a TPase coding sequence flanked by terminal inverted repeats (TIRs) (*43*). Thus, *AtMu1* provides a unique system to directly test whether the TPase itself possesses intrinsic transcriptional regulatory activity. The Arabidopsis Col-0 genome contains three closely related *AtMu1* copies (*AtMu1a*-*c*, hereafter, the *AtMu1* family) (*45*). Whole-genome resequencing data from inbred *ddm1* mutants confirmed that all three retain transposition activity, with *AtMu1a* showing the highest activity (Fig. 4A, fig. S7A). To explore potential anti-silencing activity, we independently transformed each *AtMu1* copy into WT plants and performed methylome analysis. Among the three copies, *AtMu1c* transgenic plants exhibited pronounced DNA hypomethylation at the termini of *AtMu1* family members (Fig. 4B, fig. S7B). This hypomethylation was highly specific among *AtMu* transposons, affecting almost exclusively these three *AtMu1* family members. Consistent with this epigenetic change, endogenous *AtMu1a* and *AtMu1b* were transcriptionally activated in *AtMu1c* transgenic plants (Fig. 4C). Because *AtMu1c* encodes no accessory protein, these results indicate that the AtMu1c-TPase is sufficient to induce transcriptional activation of cognate elements. To directly assess DNA-binding activity and identify genomic binding sites, we performed ChIP-seq analysis using AtMu1c-3xFLAG transgenic plants. Three independent experiments reproducibly detected sharp enrichment peaks at the TIRs of endogenous *AtMu1* copies (Fig. 4D). In addition, our ChIP-seq analysis revealed that AtMu1c-TPase bound genomic regions outside the *AtMu1* family, with *BRODYAGA2* transposons accounting for approximately one third of these additional targets (Fig. 4E, fig. S8A and B). *BRODYAGA2* represents the third most abundant TIR-type transposon family in the Arabidopsis genome (*46*). Phylogenetic analysis suggests that they are non-autonomous elements related to the *AtMu1* family (fig. S8C). Notably, the *AtMu1* family and *BRODYAGA2* shared putative target motifs of AtMu1c-TPase (Fig. 4F), and AtMu1c-TPase bound *BRODYAGA2* elements *in vivo* (Fig. 4G). Targeted *BRODYAGA2* copies also exhibited DNA hypomethylation in *AtMu1c* transgenic plants, although the effect was weaker than at *AtMu1* copies (fig. S8D).

**Fig. 4.**
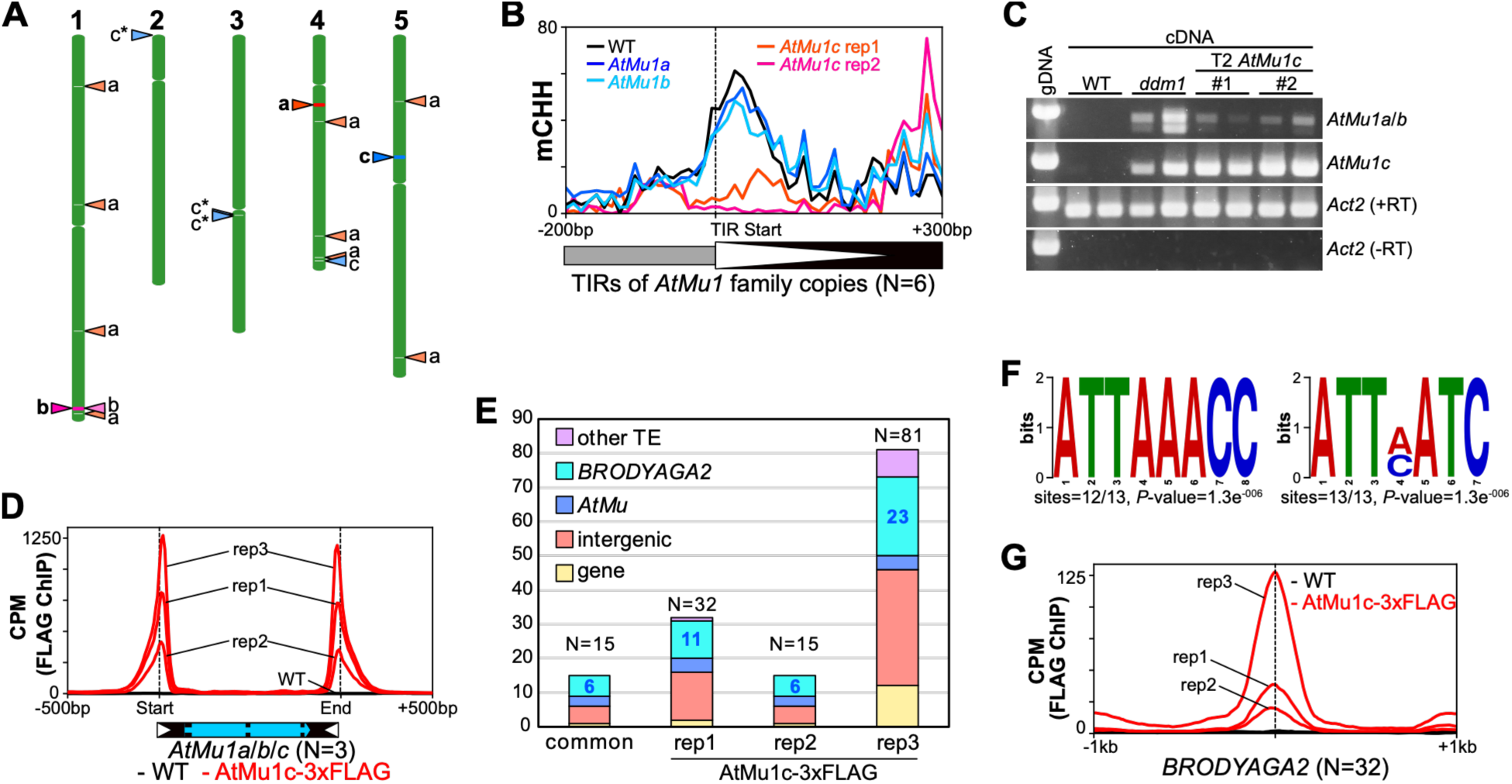
Intrinsic transcriptional regulatory activity of AtMu1c-TPase. (**A**) *De novo* insertions of *AtMu1* family transposons detected in self-pollinated *ddm1* mutants. Green boxes represent chromosomes, and triangles on the left side indicate the position of endogenous *AtMu1* family elements in the WT background. For *AtMu1c*, one non-reference insertion was detected, whose flanking sequence multiply mapped to three genomic regions with identical sequence and is indicated as “c*” on the left side of chromosomes. Non-reference *de novo* insertions are shown as triangles on the right sides of the chromosomes. (**B**) Metaplot of DNA methylation levels at TIR regions of *AtMu1* family transposons in WT and *AtMu1* transgenic plants. The vertical dashed line indicates the start point of TIRs, and 200bp upstream and 300bp downstream regions are shown. Black and gray boxes represent the *AtMu1* transposon body and flanking region, respectively. White triangle indicates TIR. (**C**) Expression of *AtMu1* TPases detected by RT-PCR. TPases of *AtMu1a* and *AtMu1b* transcripts could not be distinguished due to their high sequence similarity. Expression of AtMu1c-TPase in *AtMu1c* transgenic plants includes both transgenic and endogenous transcripts. cDNA from *ddm1* mutants was used as a positive control. (**D**) Metaplot of ChIP-seq signals (CPM) for AtMu1c-3xFLAG across *AtMu1* family transposons (N=3) including 500 bp of flanking sequence. Vertical dashed lines indicate the termini of *AtMu1* transposons. (**E**) Genomic features of AtMu1c-TPase binding targets identified by ChIP-seq analysis. (**F**) Putative target motifs of AtMu1c-TPase identified from common peaks of ChIP-seq analysis within *AtMu1* and *BRODYAGA2* transposons. (**G**) Metaplot of ChIP-seq signals (CPM) for AtMu1c-3xFLAG centered on MACS3 peaks summits of rep3 within *BRODYAGA2* transposons. Vertical dashed lines indicate the centers of the ChIP-seq peaks.

Together, these results demonstrate that AtMu1c-TPase intrinsically possesses core properties of TFs, including sequence-specific DNA-binding and the ability to induce transcriptional activation. The abundance of non-autonomous elements further suggests that a large latent *cis*-regulatory network of TPase-binding sites is pre-established in the Arabidopsis genome, enabling TPase-mediated *trans*-activation.

### One-step evolution of TFs from TPases

Domestication of TPases into TFs has been proposed to occur through fusion with host-encoded regulatory domains such as KRAB that supply the transcriptional activity (*12*). The intrinsic transcriptional regulatory activity in TPases raises the possibility of a more direct route to domestication. To test this hypothesis, we searched for high-confidence host-encoded genes annotated in RefSeq across eukaryotes that retain structural similarity to *MuDR* TPases and show signatures of functional domestication, including expression evidence, loss of transposon structural features, and low copy number. This analysis yielded 309 high-confidence TPase-derived genes spanning 38 independent domestication events, including previously characterized TPase-derived TF families such as *FAR1* and *MUSTANG* (*MUG*) (Fig. 5A, Data S1). Consistent with previous evolutionary analysis, these TF families represent multiple independent domestication events (*47*, *48*). TPase-derived TFs are expected to retain DNA-binding and transcriptional regulatory domains while losing the catalytic activity required for transposition, which in *MuDR* TPases relies on the deeply conserved DDE triad (Fig. 5B to E). Domesticated TPase-derived proteins identified here recurrently showed disruption of the DDE catalytic motif, either through substitution of one or more catalytic residues or through changes predicted to impair their spatial coordination (Fig. 5C to E). For example, *MUG* TFs carry a degenerated DDE triad, whereas the *FAR1* TF retains the three residues but harbors additional changes that are predicted to disrupt the catalytic geometry (Fig. 5F). The recurrent loss of catalytic integrity among domesticated *MuDR* TPases, together with the transcriptional regulatory activity of AtMu1c-TPase, support a one-step model in which TFs can emerge directly from TPases through the loss of transposition activity while retaining pre-existing DNA-binding and transcriptional regulatory capacity (Fig. 5G).

**Fig. 5.**
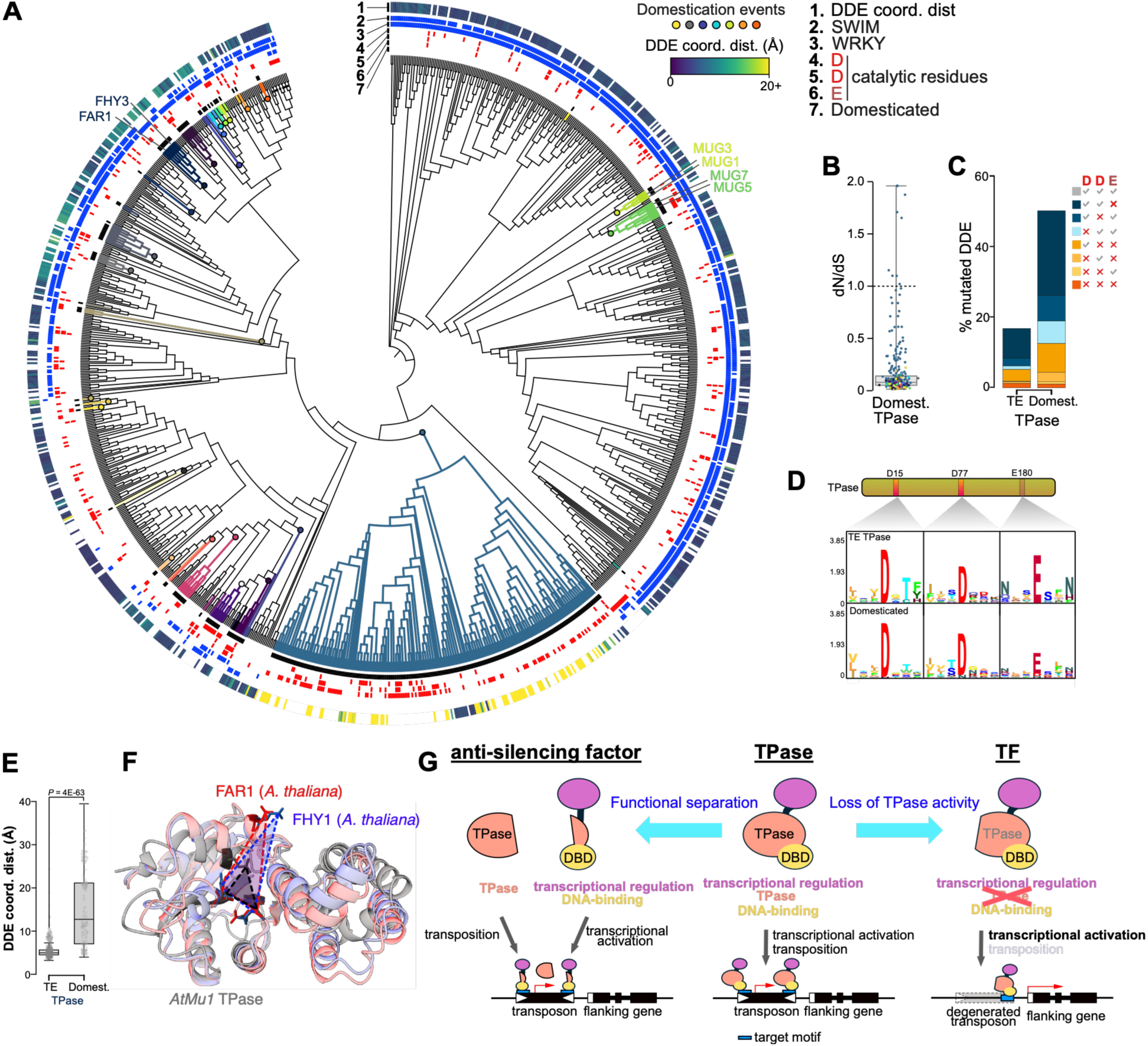
Recurrent evolution of TPases into TFs. (**A**) Cladogram of the evolutionary history of the DDE-containing domain from TPases and TPase-like proteins. Independent domestication events are marked as colored branches and shown in black (track 7). Presence of the additional SWIM and WRKY domains on the full-length proteins is shown in blue (track 2 and 3, respectively). Proteins with any of the catalytic residues (D, D and E) mutated are highlighted in red (track 4 to 6). For domains with an intact DDE, the coordination distance between the three residues is shown as a heatmap. (**B**) *dN/dS* ratios of domesticated TPases. (**C**) Percentage of *MuDR* TPases and domesticated TPases with substitutions in the catalytic residues D, D and E. (**D**) Amino acid sequence logo surrounding the three catalytic residues for both TPases and TPase-like proteins. (**E**) Distribution of coordination distances for TPases and domesticated TPase-like proteins. Only proteins with an intact DDE are included. Statistical significance of differences was calculated using a two-sided Mann-Whitney U test. (**F**) Illustrative example of the 3D structure of the AtMu1c, FAR1 and FHY3 DDE-containing domains. The residues forming the DDE (DDD for FHY3) and the interatomic distance among them is highlighted. (**G**) Proposed model for the evolution of TFs from TPase.

## Discussion

In this study, we show that sequence-specific transcriptional regulatory activity can be encoded by a TPase. Through analyses of the *Spm* and *AtMu1* families, we demonstrate that TPase and TPase-derived factors can induce sequence-specific DNA hypomethylation and transcriptional activation. In this sense, transposons can function as self-contained TFs. This provides a mechanistic explanation for the convergent evolution of anti-silencing systems across unrelated transposon superfamilies (*19*-*21*, this study) and establish a framework for understanding how transposons directly participate in transcriptional control.

Our findings also provide a unifying framework for interpreting earlier observations of epigenetic changes induced by TPases, including *MuDR*, *hAT*, and *Ac* in maize and *Tam3* in snapdragon (*Antirrhinum majus*) (*49–52*). DNA hypomethylation induced by TPases of these transposons can be eventually restored once functional TPase is dissociated or segregated (*51*, *52*), closely mirroring the reversible behavior of VANC-and TnpA-mediated anti-silencing systems (*19*, *27*, Fig. 1). These parallels suggest that transcriptional regulatory activities, including anti-silencing activity, may represent a widespread and ancestral property of TPases.

Transposition is known to shape transcriptional networks by dispersing TF-binding sites throughout the genome. Our results extend this view by demonstrating that transposons themselves can autonomously establish *cis*-regulatory networks through self-binding and transcriptional activation. Transposon-mediated latent transcriptional networks may be extensive because TPases bind not only autonomous elements but also abundant non-autonomous copies, as shown for *AtMu1c* (Fig. 4). Although epigenetic silencing normally masks these interactions, transient transposon activation, particularly during epigenetic reprogramming in sexual lineages, could expose them (*53*).

Together, these observations support a parsimonious model in which the intrinsic capacity of TPases to transcriptionally regulate cognate transposons provides a pre-existing basis for their domestication as TFs, while loss of transposition activity facilitates their co-option as host transcriptional regulators (Fig. 5G). While recent work has demonstrated that TFs can evolve through fusion of host transcriptional regulatory domains with TPase-derived DBDs (*12*), our results uncover an alternative and more direct evolutionary route from TPases to TFs. Several plant TF families, including *FHY3*, *FAR1* and *MUG*, are derived from *MuDR* TPase and harbor mutations that disrupt the conserved DDE catalytic motif required for transposition (*10*, *54*). Our research expands this evolutionary pathway by identifying additional candidates of domesticated genes from *MuDR*-related TPases (Fig. 5). These observations suggest that many TPases may function as proto-TFs, carrying latent regulatory potential even before domestication. Because transposition continuously generates numerous TPase copies throughout the genome and each TPase copy already carries intrinsic transcriptional regulatory activity, every new copy represents an independent substrate for domestication through loss of catalytic activity. This model provides a mechanistic explanation for the remarkable observation that members of the same TPase-derived TF family, such as *FAR1* and *MUG*, repeatedly originated through independent domestication events during evolution (*47*, *48*).

In the case of *Spm* transposons, the TnpA appears to have evolved through functional separation from the ancestral TPase. This separation likely reduces the cytotoxic risks associated with transposition while allowing regulatory functions to diversify under relaxed evolutionary constraints. Additional changes, including loss of TIRs and remodeling of protein domains, likely contributed to the functional diversification of TPases into TFs. These evolutionary principles may be broadly applicable to TPases across the tree of life.

More than seven decades after Barbara McClintock introduced transposons as “controlling elements”, our findings provide a molecular basis for this concept by showing that novel transcriptional networks can emerge from within latent transcriptional regulatory systems comprising TPases and their cognate sequences.

**fig. S1.**
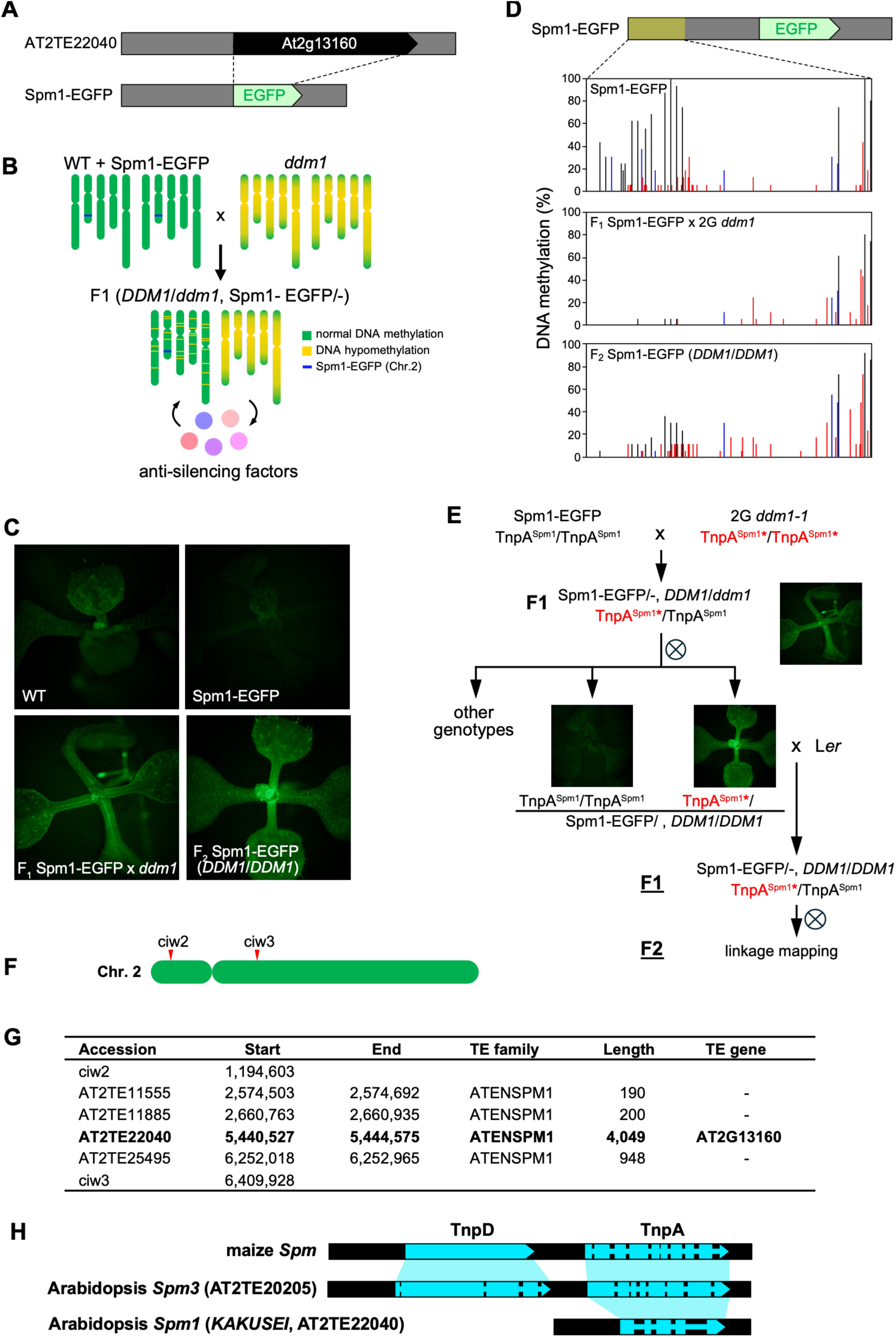
Identification of the novel anti-silencing factor TnpA^Spm1^. (**A**) Schematic representation of Spm1-EGFP reporter construct. The gene At2g13160 within AT2TE22040 was replaced with EGFP. (**B**) Genetic strategy for *trans*-activation of the Spm1-EGFP reporter via crossing with *ddm1* mutant. Hypomethylation status induced by *ddm1* mutation is epigenetically heritable even after backcrossing with WT plants. Chromosomal regions in green and yellow represent normally methylated and hypomethylated status, respectively. (**C**) EGFP fluorescence in seedlings of parental WT, Spm1-EGFP, their F1 progeny, and F2 plants homozygous for the WT *DDM1* allele. (**D**) DNA methylation status of 5’ region of the Spm1-EGFP reporter detected by bisulfite sequencing. Black, blue, and red vertical bars indicate CG, CHG, and CHH methylation, respectively. For each sample, 16 copies were sequenced. (**E**) Genetic pedigree used to generate the mapping population. TnpA^Spm1^* denotes the *ddm1*-derived epigenetically activated TnpA^Spm1^ allele. (**F**) Genomic region associated with *trans*-activation of the Spm1-EGFP reporter. ciw2 and ciw3 are markers flanking to the candidate locus. (**G**) List of *Spm1* transposons within the mapped region. Only AT2TE22040 (*KAKUSEI*) is a full-length element that encodes a protein. (**H**) Comparison of transposon structures among maize *Spm*, Arabidopsis *Spm3* (AT2TE20505), and *KAKUSEI*. Blue and black regions indicate coding and noncoding regions, respectively.

**fig. S2.**
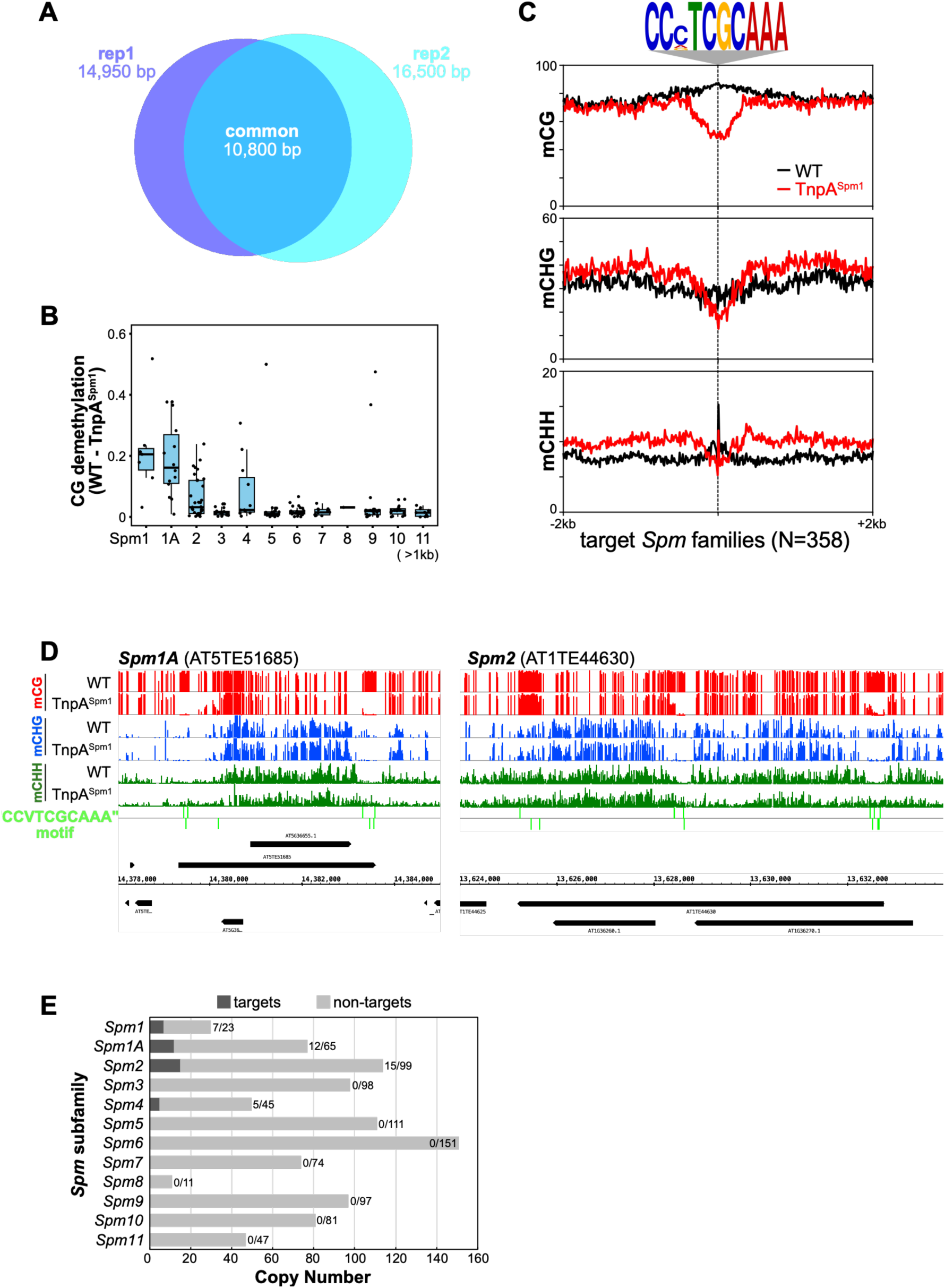
Effects of TnpA^Spm1^ on DNA methylation. (**A**) Venn diagram showing the total length of CG hypomethylated regions induced by TnpA^Spm1^ and detected in two independent replicates of methylome analysis. (**B**) Boxplot showing the decrease in CG methylation for each *Spm* copy induced by TnpA^Spm1^ across *Spm* subfamilies. Average values from two methylome datasets were calculated for *Spm* transposons longer than 1 kb. (**C**) Metaplot of DNA methylation levels in the flanking regions of target “CCVTCGCAAA” motif within target *Spm* families (*Spm1*, *Spm1A*, *Spm2*, and *Spm4*). Black and red lines indicate WT and TnpA^Spm1^, respectively. (**D**) Representative examples of DNA methylation patterns in WT and TnpA^Spm1^. (**E**) Number of hypomethylated transposons targeted by TnpA^Spm1^. Elements with at least one CG hypoDMR (50-bp bin, ΔmCG > 0.4) consistently detected in both replicates were counted as targets.

**fig. S3.**
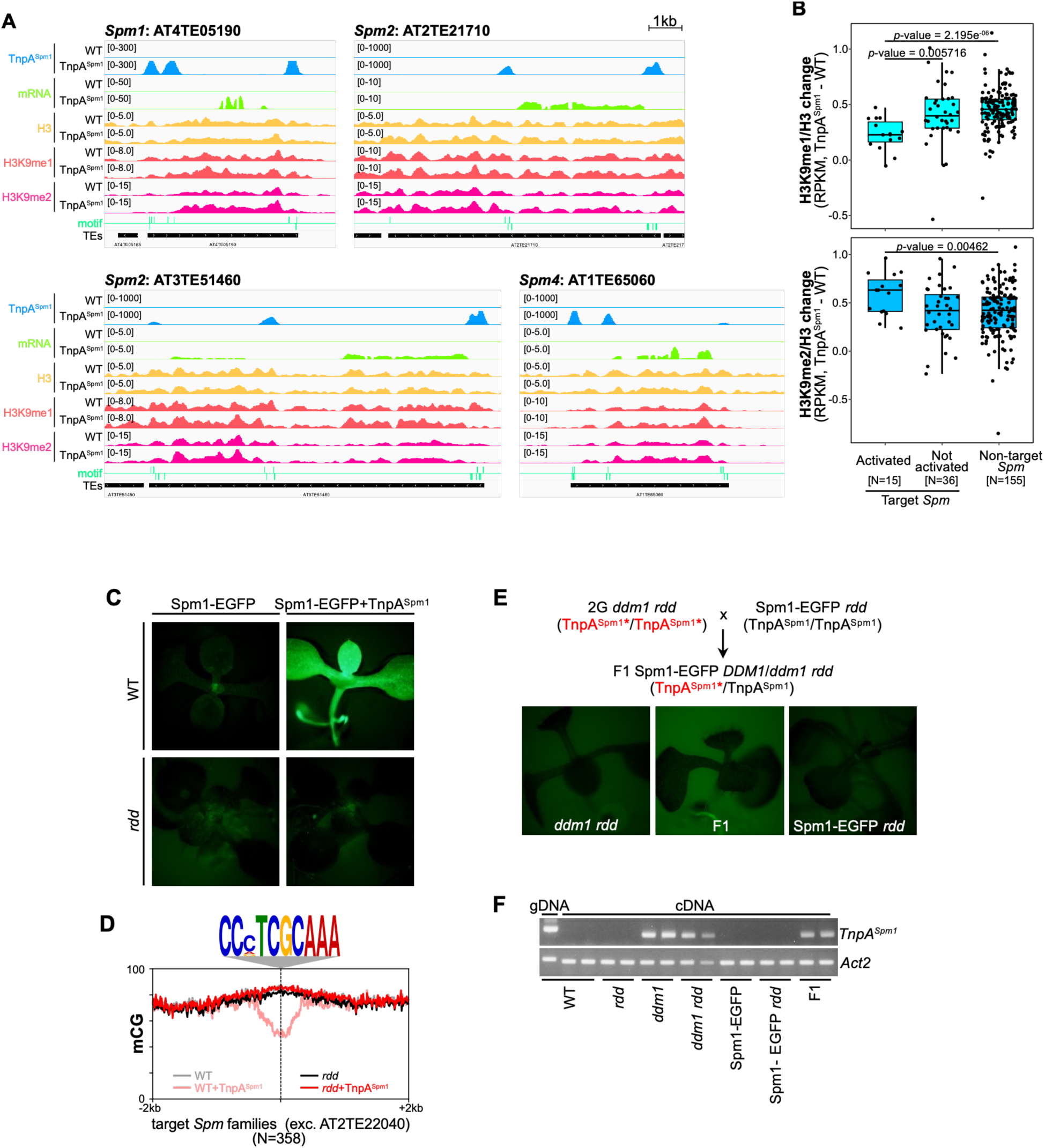
DNA demethylases are required for TnpA^Spm1^-mediated anti-silencing. (**A**) Genome browser views of representative TnpA^Spm1^ targets, showing ChIP-seq signals for TnpA^Spm1^-3xFLAG, transcription, and histone modifications (H3, H3K9me1, and H3K9me2) in WT and TnpA^Spm1^. (**B**) Boxplot showing changes in H3K9me1 (top) and H3K9me2 (bottom) levels in TnpA^Spm1^ compared to WT (TnpA^Spm1^ - WT). Signals were calculated in *Spm* transposons harboring TE genes, and normalized to H3 (RPKM, H3K9me/H3). “Target *Spm*” refers to *Spm1*, *1A*, *2*, and *4*. “Non-target *Spm*” refers to other *Spm* subfamilies. “Target *Spm*” is further divided into “Active” (log2 fold change >1.5 in transcript level [RPKM+1] in at least one of two RNA-seq replicates) and “Not activated”. The P-values are based on a Kolmogorov-Smirnov test. (**C**) EGFP fluorescence in WT and *rdd* backgrounds expressing TnpA^Spm1^, observed as in Fig. 1B. (**D**) Patterns of CG methylation in 2-kb flanking regions of “CCVTCGCAAA” motifs. Black and red lines indicate non-transgenic and TnpA^Spm1^ plants, respectively. Light and dark shades represent WT and *rdd* backgrounds, respectively. (**E**) *Trans*-activation of Spm1-EGFP reporter in *rdd* background by genetic crossing, as in fig. S1C. (**F**) RT-PCR showing TnpA^Spm1^ expression in plants used in (E) with positive (*ddm1*) and negative (WT, *rdd*, Spm1-EGFP) controls. *Act2* was used as control.

**fig S4.**
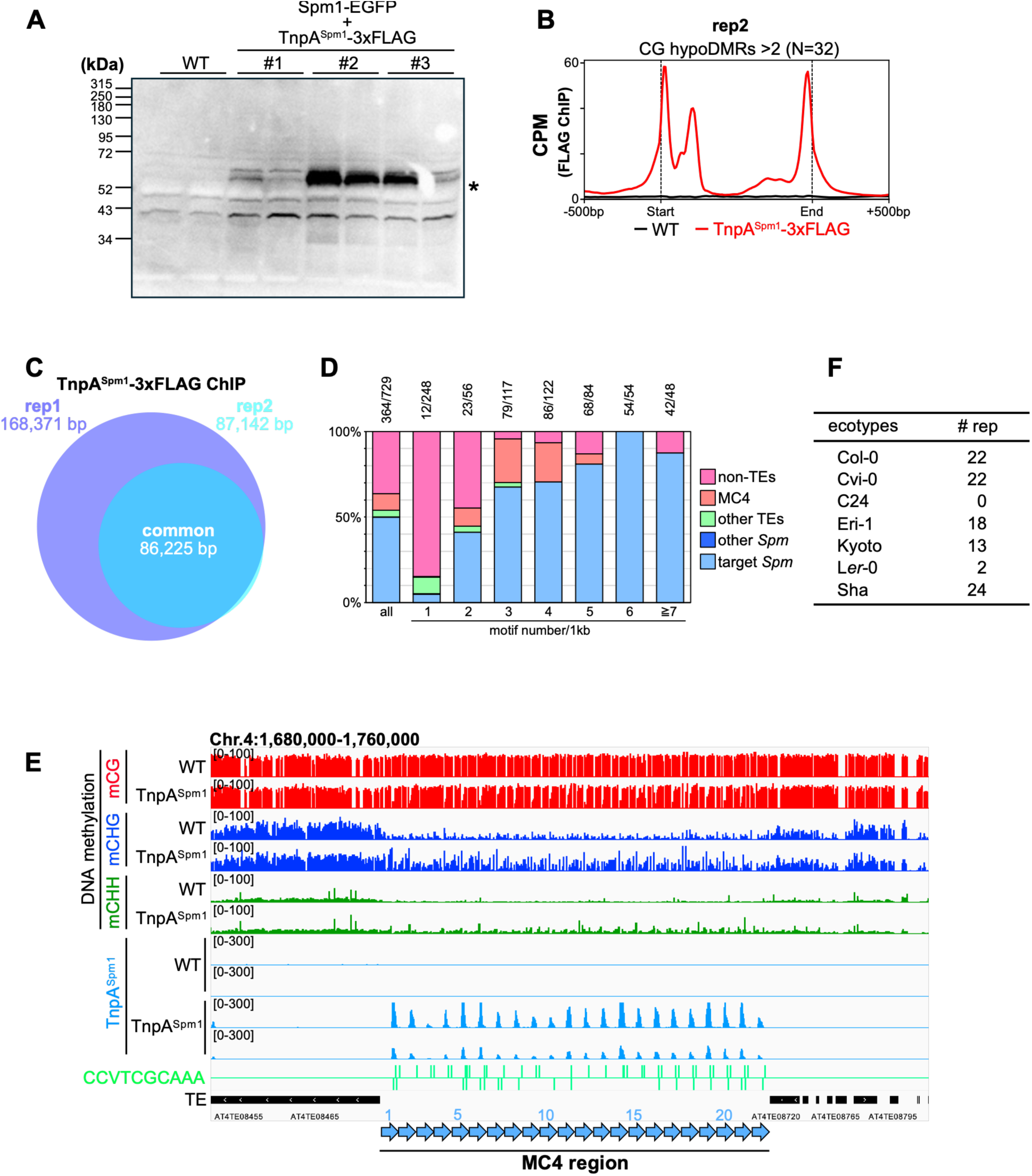
Identification of TnpA^Spm1^ target sites by ChIP-seq analysis. (**A**) Immunoblot detecting TnpA^Spm1^-3xFLAG expression. The asterisk indicates the expected signal for the tagged protein. (**B**) Metaplot of ChIP-seq signals (CPM; counts per million reads) for TnpA^Spm1^-3xFLAG across *Spm* transposons containing ≥2 CG hypoDMRs (N=32). Black and red lines represent WT and TnpA^Spm1^-3xFLAG, respectively. This dataset corresponds to an independent replicate of Fig. 1E. (**C**) Venn diagram showing overlap in TnpA^Spm1^-binding regions identified in two independent ChIP-seq experiments. (**D**) Distribution of the “CCVTCGCAAA” motifs across different genomic components, grouped by motif density (motif number per 1kb). Total motifs counts and counts within target *Spm* transposons are shown above the bars. (**E**) Genome browser view of the MC4 region showing DNA methylation status, ChIP-seq signals for TnpA^Spm1^-3xFLAG in WT and TnpA^Spm1^ transgenic plants, and target motif distribution. (**F**) Numbers of repeat units in the MC4 region among different Arabidopsis ecotypes.

**fig S5.**
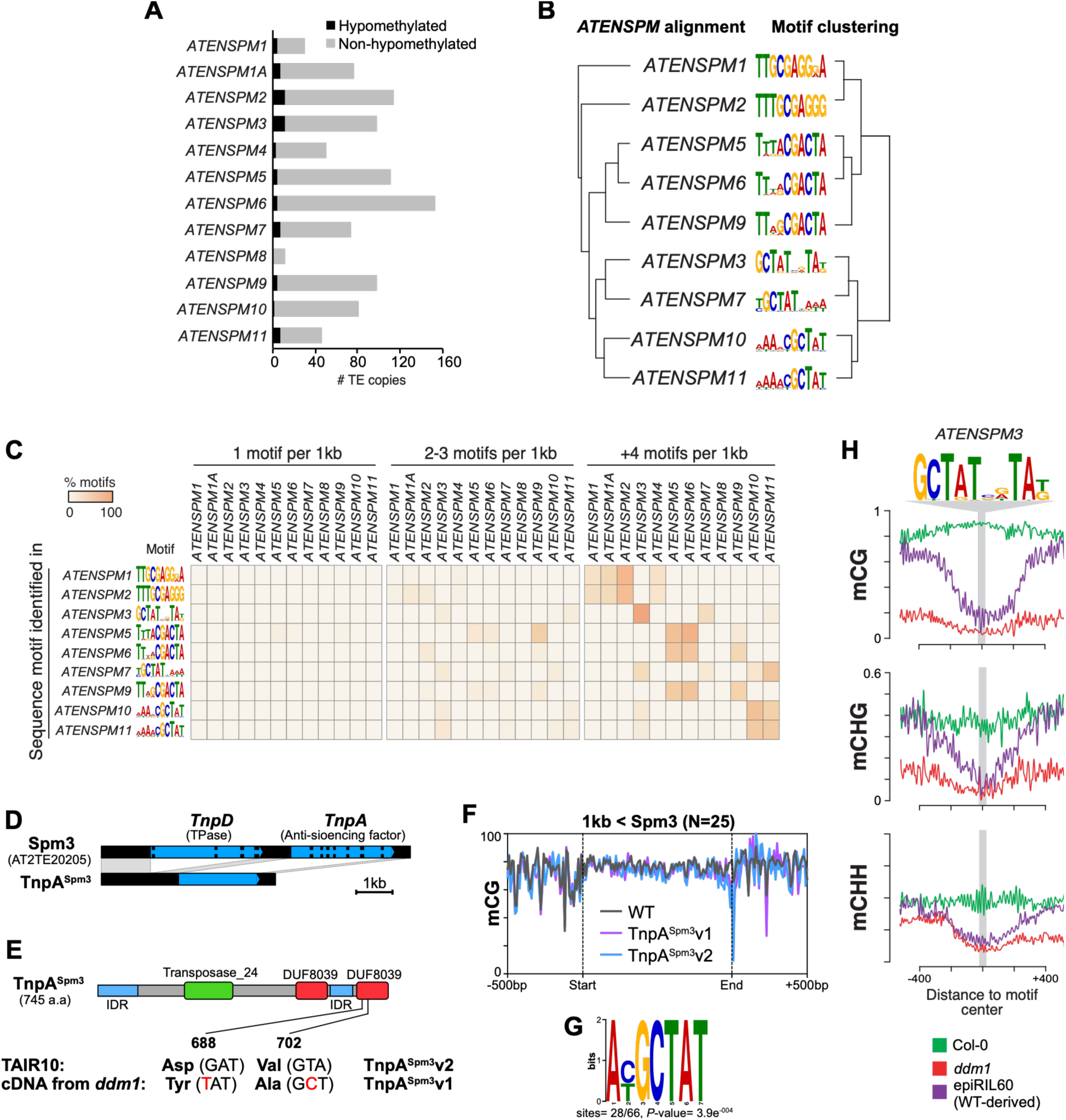
Comprehensive identification of TnpAs in *Spm* subfamilies. (**A**) Numbers of *trans*-hypomethylated *Spm* copies detected in epiRILs for each *Spm* family. (**B**) Phylogenetic relationship of *Spm* families (left, based on RepBase consensus sequences) and similarity-based clustering of their putative target motif sequences (right). (**C**) Heatmap showing the proportion of transposon copies in each *Spm* family classified according to motif density. Motif density categories were defined as low (1 motif/1 kb), medium (2–3 motifs/1 kb), and high (+4 motifs/1 kb). (**D**) Schematic representation of the autonomous *Spm3* copy AT2TE20205 and the TnpA^Spm3^ construct. Blue and black regions indicate coding and noncoding regions, respectively. (**E**) Schematic diagram of the protein structure of TnpA^Spm3^. The cDNA clones obtained from *ddm1-1* harbor two nonsynonymous mutations (D688Y and V702A) relative to the TAIR10 reference sequence. Two TnpA^Spm3^ constructs were generated: one containing the *ddm1* cDNA sequence (TnpA^Spm3^v1) and the other containing the TAIR10 sequence (TnpA^Spm3^v2). (**F**) Metaplot showing CG methylation levels across *Spm3* transposons (>1 kb, N=25) and their flanking regions. Dashed vertical lines indicate the start and end positions of *Spm3* transposons. DNA methylation levels in WT, TnpA^Spm3^v1, and TnpA^Spm3^v2 are shown with black, purple, and blue lines, respectively. (**G**) Putative target motif of TnpA^Spm3^ predicted by DREME using CG hypoDMRs. (**H**) Metaplot of DNA methylation in 500bp flanking regions of putative target motif of TnpA^Spm3^ in WT (green), *ddm1* (red), and WT-derived regions of epiRIL60 (purple).

**fig S6.**
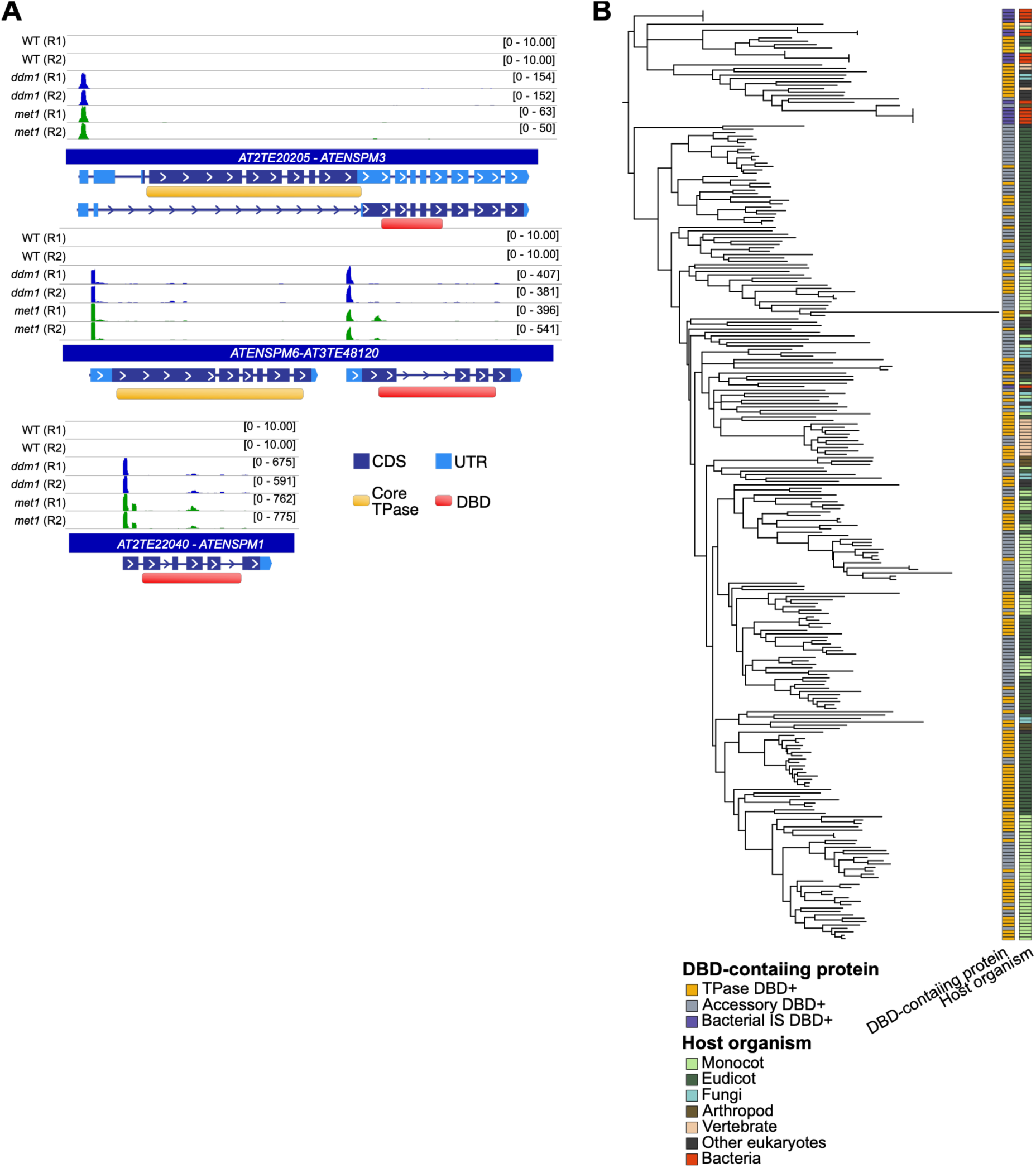
Alternative transcriptional organization of *Spm*-derived DNA-binding domain (DBD) proteins. (**A**) Cap Analysis of Gene Expression sequencing (CAGE-seq) profiles from wild-type (WT), *ddm1*, and *met1* plants across representative Arabidopsis *Spm* elements. Gene models are shown below each locus, with coding sequences (CDS) in dark blue and untranslated regions (UTRs) in light blue. The positions of the conserved TPase catalytic core and DNA-binding domain (DBD) are indicated in orange and red, respectively. Top, AT2TE20205-ATENSPM3, in which alternative splicing generates transcripts encoding either the TPase catalytic core or the DBD. Middle, ATENSPM6-AT3TE48120, in which TPase-and DBD-coding sequences are transcribed from separate genes within the same locus. Bottom, AT2TE22040-ATENSPM1 (*KAKUSEI*), encoding a single DBD-containing transcript. CAGE-seq signals are shown for two biological replicates of WT, *ddm1*, and *met1* plants. (**B**) Phylogenetic tree of protein sequences spanning DBD domains identified in *Spm* TPases (yellow) or accessory proteins (grey). Bacterial tri-helical HTH DBDs from *E. coli* insertion sequences (IS, violet) were included to root the tree. Sequences were aligned with MAFFT E-INS-i and the tree inferred with IQ-TREE3 using 1,000 bootstrap replicates; branches with <50% support were collapsed. The organisms encoding the analyzed proteins are shown.

**fig S7.**
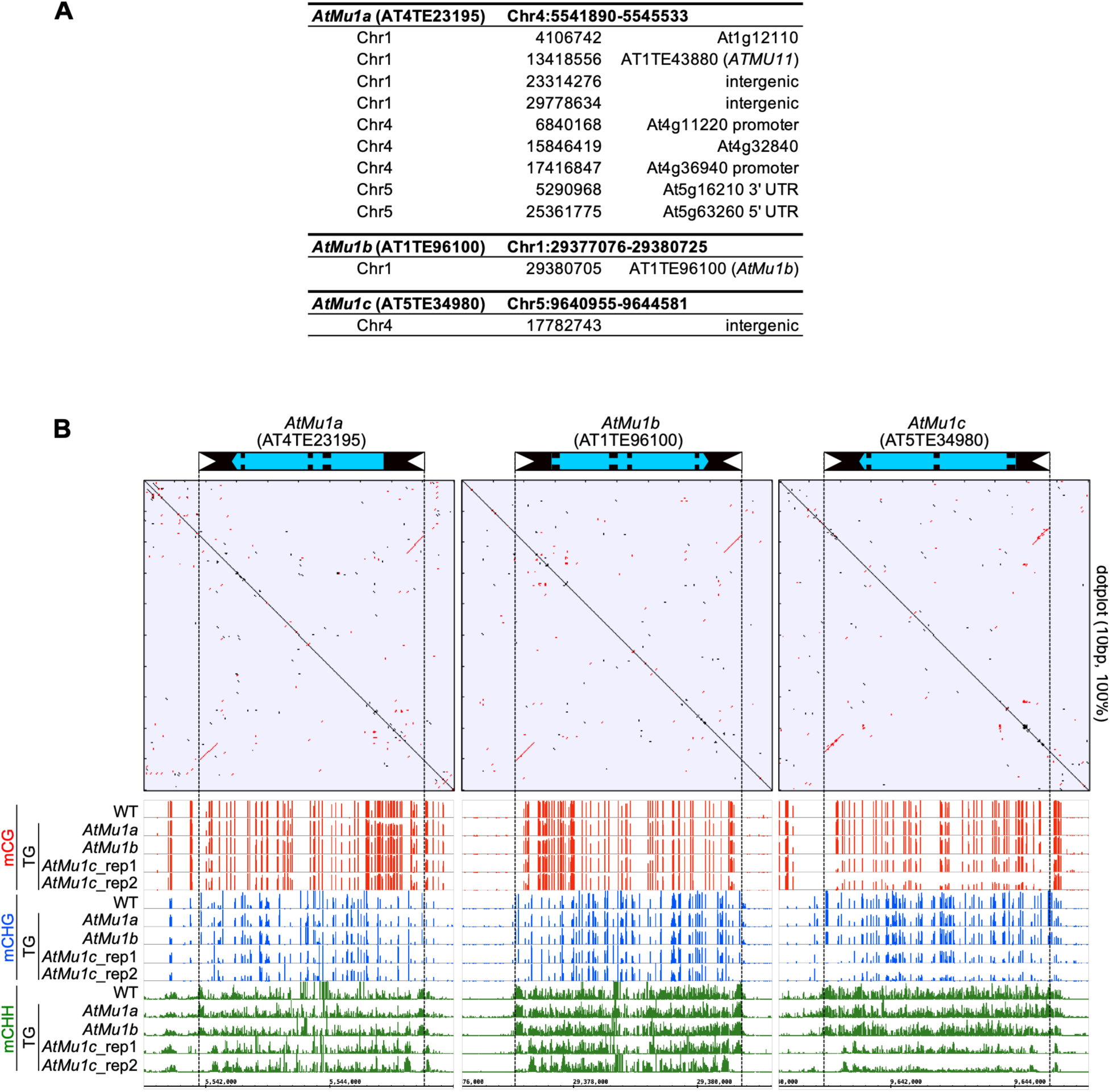
*AtMu1c* transposon exhibits both anti-silencing and transposition activities. (**A**) Genomic positions and properties of *de novo* insertions of the *AtMu1* family. Chromosome, inserted position, and characteristics of the insertion sites are listed. (**B**) Schematic structures (top), dot-plot analyses (middle), and DNA methylation profiles (bottom) of each *AtMu1* family transposon in *AtMu1* transgenic plants. In the schematic structures, white triangles indicate TIRs, and blue and black regions represent coding and noncoding regions, respectively. Dot-plot analyses were performed using 10-bp windows with 100% sequence identity. Black and red lines represent forward and reverse strand matches, respectively. Genomic coordinates of *AtMu1* family transposons are corrected based on the positions of the TIRs and TSDs. For DNA methylation profiles, red, blue, and green lines indicate the proportions of methylated cytosines in CG, CHG, and CHH (H: A, T, or C) contexts, respectively.

**fig S8.**
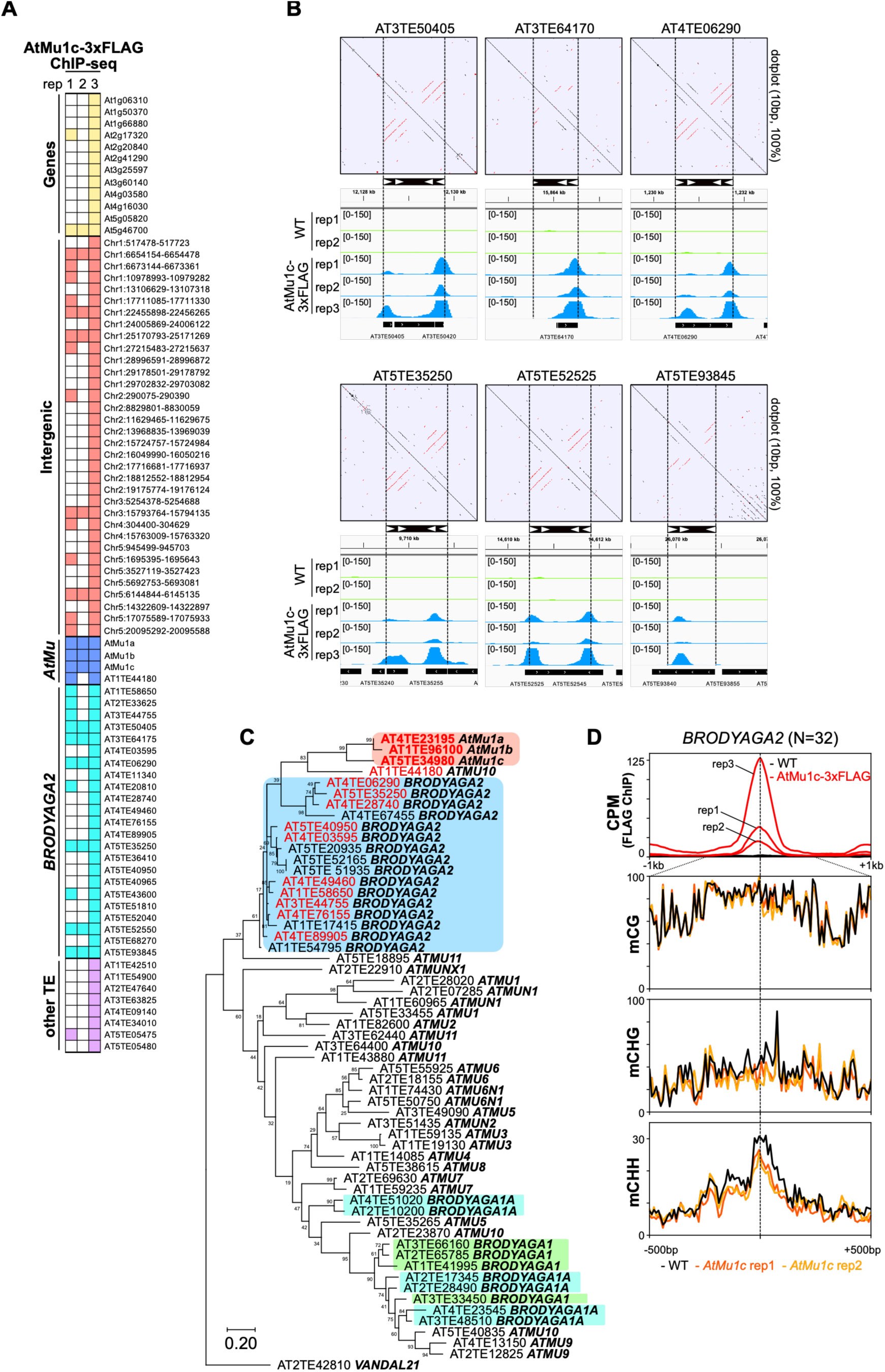
Pre-establishment of *cis*-regulatory network integrated by AtMu1c-TPase. (**A**) Heatmap showing AtMu1c-TPase binding targets across three independent ChIP-seq experiments. Colored boxes indicate ChIP-seq signals detected by MACS3 in each ChIP-seq experiment. (**B**) Genome browser views of ChIP-seq signals for AtMu1c-3xFLAG within *BRODYAGA2* transposons reproducibly detected in three independent ChIP-seq experiments. Dot-plot analyses and schematic structures of *BRODYAGA2* transposons are also shown. (**C**) Phylogenetic tree of full-length *AtMu* and *BRODYAGA2* transposons. For each copy, 120-bp sequences from 5’ terminal regions were used. Copies analyzed are listed in Table S3 (*AtMu*) and S4 (*BRODYAGA2*). Transposon copies with ChIP-seq signal of AtMu1c-3xFLAG in at least one ChIP experiment are shown in red. (**D**) Metaplot of ChIP-seq signals (CPM) (same as Fig. 4G) and DNA methylation levels for AtMu1c-TPase -binding *BRODYAGA2* transposons (N=32). Vertical dashed lines indicate the centers of the ChIP-seq peaks. DNA methylation levels in WT and two independent *AtMu1c* transgenic plants are shown.

**Table S1.**
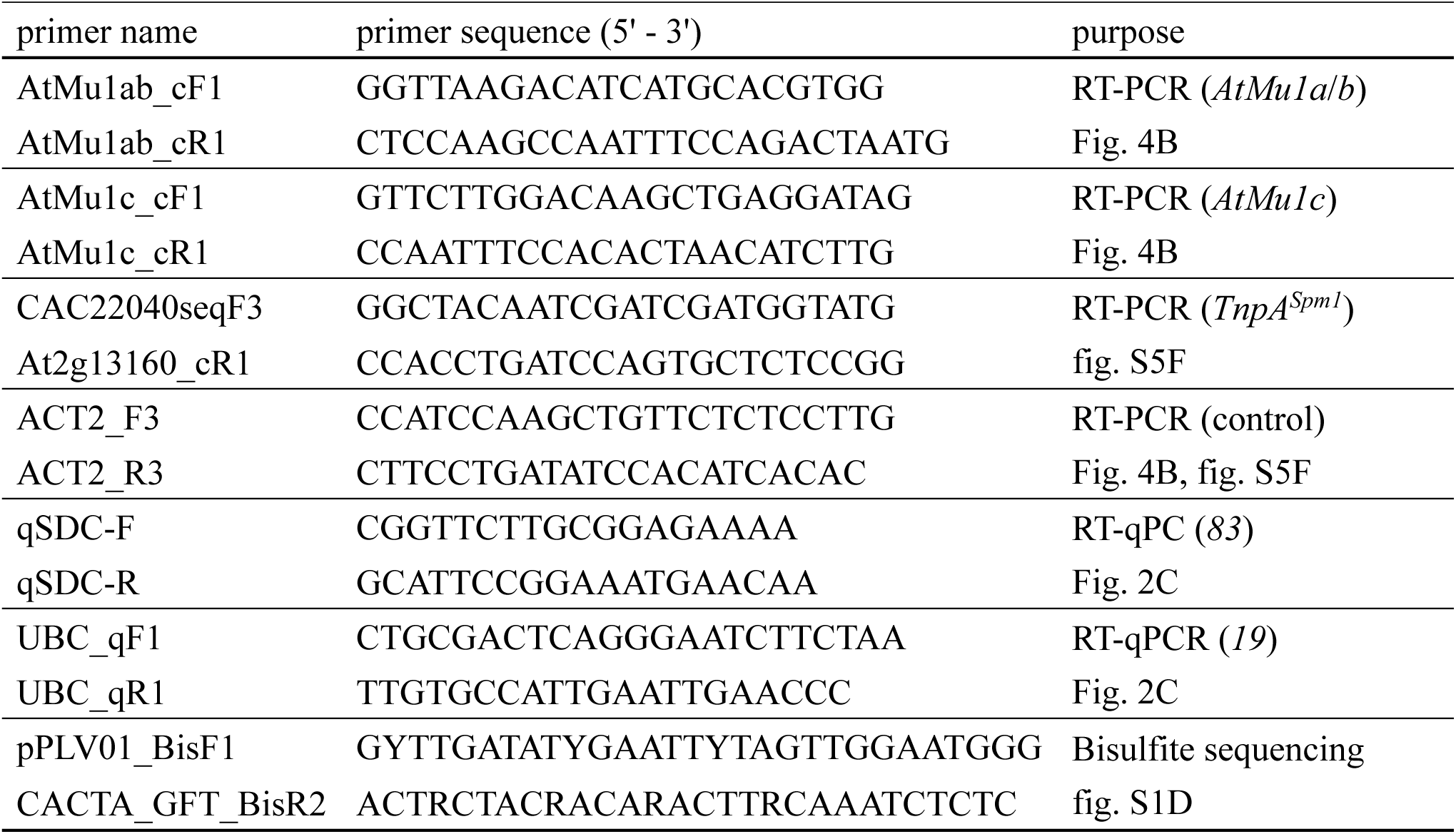
Primers used in this study.

**Table S2.** List of full-length *Spm* transposons.

| TE accession | TE family | Chr | start | end | length | direction | TSD | TAIR10 annotation |
| --- | --- | --- | --- | --- | --- | --- | --- | --- |
| AT1TE38940 | ATENSPM1A | 1 | 12016041 | 12020186 | 4146 | - | AAA | Chr1:12016038-12020227 |
| AT1TE41230 | ATENSPM1A | 1 | 12626457 | 12630547 | 4091 | + | ATT | Chr1:12626457-12630547 |
| AT1TE42210 | ATENSPM3 | 1 | 12942847 | 12947011 | 4165 | + | CTA | Chr1:12942825-12947010 |
| AT1TE45020 | ATENSPM5 | 1 | 13734686 | 13743202 | 8517 | - | TAT | Chr1:13734680-13743201 |
| AT1TE52590 | ATENSPM2 | 1 | 15950604 | 15958318 | 7715 | - | ACA | Chr1:15950604-15958318 |
| AT1TE60180 | ATENSPM9 | 1 | 18154506 | 18163468 | 8963 | + | ACA | Chr1:18154505-18163494 |
| AT1TE65060 | ATENSPM4 | 1 | 19675548 | 19680346 | 4799 | + | TAC | Chr1:19675549-19680353 |
| AT2TE16310 | ATENSPM7 | 2 | 3785520 | 3794054 | 8535 | - | ACG | Chr2:3785520-3794054 |
| AT2TE17600 | ATENSPM6 | 2 | 4162311 | 4169555 | 7245 | + | CGA | Chr2:4162311-4169555 |
| AT2TE18415 | ATENSPM3 | 2 | 4385856 | 4394311 | 8456 | - | TTT | Chr2:4385856-4394311 |
| AT2TE20205 | ATENSPM3 | 2 | 4900803 | 4909281 | 8479 | + | ATA | Chr2:4900803-4909281 |
| AT2TE20255 | ATENSPM5 | 2 | 4925182 | 4933932 | 8751 | - | TTA | Chr2:4925171-4933931 |
| AT2TE21710 | ATENSPM2 | 2 | 5338034 | 5346286 | 8253 | - | ATA | Chr2:5338033-5346285 |
| AT2TE22040 | ATENSPM1 | 2 | 5440527 | 5444575 | 4049 | + | TCT | Chr2:5440527-5444575 |
| AT2TE23705 | ATENSPM4 | 2 | 5817351 | 5822205 | 4855 | + | AGA | Chr2:5817335-5822205 |
| AT2TE24525 | ATENSPM1A | 2 | 6027027 | 6031165 | 4139 | - | TAG | Chr2:6027027-6031164 |
| AT2TE63800 | ATENSPM1 | 2 | 14407894 | 14412410 | 4517 | + | AGA | Chr2:14407884-14412426 |
| AT3TE47975 | ATENSPM11 | 3 | 11513763 | 11521906 | 8144 | + | GAT | Chr3:11513764-11521912 |
| AT3TE48120 | ATENSPM6 | 3 | 11560702 | 11569513 | 8812 | - | ATA | Chr3:11560702-11569513 |
| AT3TE51460 | ATENSPM2 | 3 | 12461860 | 12472012 | 10153 | - | ATA | Chr3:12461860-12472012 |
| AT3TE53045 | ATENSPM10 | 3 | 12919569 | 12927696 | 8128 | - | AGAT/AGA | Chr3:12919569-12927696 |
| AT3TE69350 | ATENSPM9 | 3 | 17106173 | 17115098 | 8926 | + | ACT | Chr3:17106151-17115098 |
| AT4TE05190 | ATENSPM1 | 4 | 1016802 | 1021355 | 4554 | + | CAT | Chr4:1016803-1021368 |
| AT4TE09085 | ATENSPM5 | 4 | 1838260 | 1846903 | 8644 | + | TAC | Chr4:1838260-1846903 |
| AT4TE10350 | ATENSPM1 | 4 | 2208276 | 2212250 | 3975 | - | GAA | Chr4:2208276-2212249 |
| AT5TE39425 | ATENSPM11 | 5 | 10846170 | 10853863 | 7694 | + | TTG/ATG | Chr5:10846170-10853863 |
| AT5TE41980 | ATENSPM10 | 5 | 11623257 | 11631509 | 8253 | - | GAG | Chr5:11623257-11631509 |
| AT5TE65540 | ATENSPM5 | 5 | 18193416 | 18202052 | 8637 | - | TAT | Chr5:18193410-18202051 |

**Table S3.**
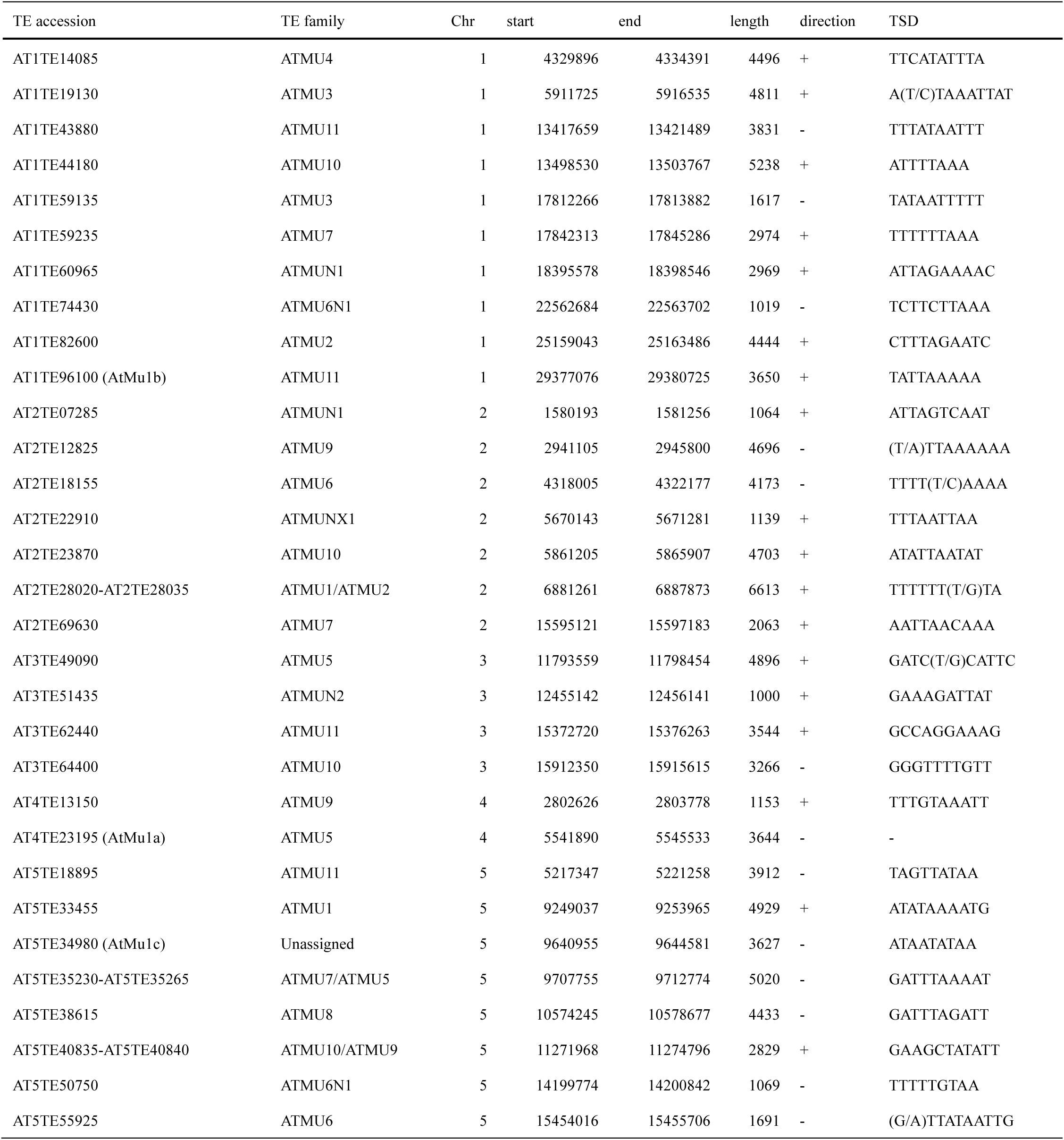
List of *AtMu* transposons analyzed in this study.

**Table S4.**
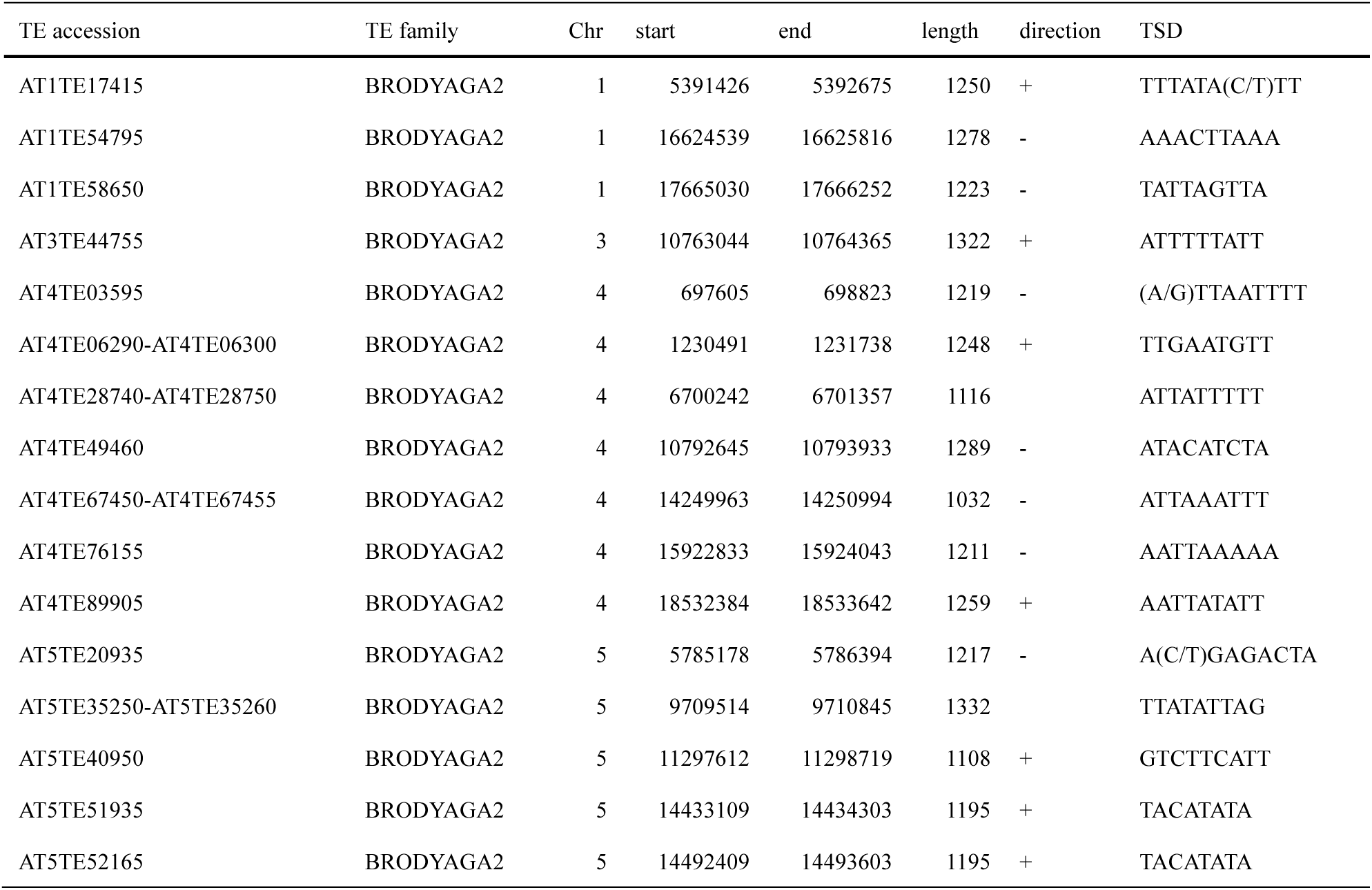
List of *BRODYAGA2* transposons analyzed in this study.

## Materials and Methods

### Plant Materials

Isolation of *ddm1-1* was reported previously (*55*). T-DNA insertion mutants of *drm1-2* (SALK_031705), *drm2-2* (SALK_150863), *cmt3-11t*(SALK_148381), *ros1-4* (SALK_045303), *dml2* (SALK_141248), *dml3-2* (SALK_056440), and *sdc-1* (SALK_017593) were obtained from ABRC (*56*). All plants used in this experiment are Col-0 background, except for that L*er* was used for linkage mapping.

### Construction and transformation

For Spm1-EGFP construct, the noncoding regions of *KAKUSEI* and EGFP sequence were amplified by PCR and assembled into pPLV01 vector digested with *Hpa* I and *Eco*53kI using NEBuilder HiFi DNA Assembly (NEB). TnpA^Spm1^ and *AtMu1* copies were amplified by PCR using genomic DNA as a template and assembled into *Sma* I-digested pGreenII-0179 using NEBuilder. For the TnpA^Spm3^v1 construct, the noncoding regions of AT2TE20205 were amplified and assembled together with the coding sequence of TnpA^Spm3^, which was amplified from cDNA synthesized using RNA extracted from *ddm1* mutants, into *Sma* I-digested pGreenII. The TnpA^Spm3^v2 construct was generated by PCR-based site-directed mutagenesis using the TnpA^Spm3^v1 construct as a template. Constructs were transformed into WT (Spm1-EGFP, TnpA^Spm3^, and *AtMu1* constructs) or Spm1-EGFP (TnpA^Spm1^ and TnpA^Spm1^-3xFLAG) plants by floral dip method using *Agrobacterium tumefaciens* strain GV3101::pMP90 (*57*).

### Linkage mapping of *TnpA^Spm1^*

To generate mapping population, Spm1-EGFP and the 2G *ddm1* mutant were crossed, and resulting F1 plants were self-pollinated. From the F2 population, EGFP-positive individuals homozygous for the WT *DDM1* allele were selected and used as the parental line for linkage analysis. This line was crossed with L*er*, and F2 seeds were sown on MS medium supplemented with phosphinothricin (for selection of Spm1-EGFP transgene). Both EGFP-positive and EGFP-negative segregants were collected and used for mapping.

### DNA methylation analysis

Genomic DNA was isolated from mature leaves using Nucleon Phytopure (GE Healthcare). For whole-genome methylome analysis, 500 ng of genomic DNA was fragmented with Covaris system, and DNA fragments of 300-450 bp were size-selected by gel electrophoresis and purified using FastGene Gel/PCR Extraction Kit (Nippon Genetics). Purified DNA was used for library construction with either KAPA HyperPrep Kit (KAPA Biosystems) or NEBNext Enzymatic Methyl-seq Kit (NEB). For library prepared with KAPA HyperPrep Kit, unmethylated cytosines were converted to uracil using MethylCode Bisulfite Conversion Kit (Applied Biosystems), followed by PCR amplification with KAPA HiFi HotStart Uracil+ ReadyMix (KAPA Biosystems). Amplified libraries were purified with AMPure XP beads (Beckman Coulter). Libraries were sequenced by Macrogen Japan or Novogene Japan.

Locus-specific Bisulfite sequencing for Spm1-EGFP was performed as previously described (*35*). Primers used for PCR amplification were listed in Table S1.

### RNA expression analysis

Total RNA was extracted from mature leaves using Trizol reagent (Thermo Fisher Scientific) and treated with DNase I (Invitrogen). For RNA-seq analysis, 600 ng total RNA was used for library preparation with KAPA mRNA HyperPrep Kit (Roche) according to the manufacturer’s instructions. For indexing and library amplification, KAPA Unique Dual Primer Kit and KAPA Universal Adapter (Roche) were used, respectively. Libraries were sequenced by Macrogen Japan or Novogene Japan.

For RT-PCR and real-time quantitative PCR (RT-qPOCR), 1 µg total RNA was used for cDNA synthesis using ReverTra Ace qPCR RT Master Mix (TOYOBO). One µL of 10-fold diluted cDNA was used as a template for RT-PCR in 10 µL reaction mix using EmeraldAmp MAX PCR Master Mix (TaKaRa). RT-qPCR was performed with PowerUp SYBR Green Master Mix (Thermo Fisher Scientific) on StepOne Real-Time PCR Systems. Primer sequences used for RT-PCR and RT-qPCR are listed in Table S1.

### eChIP-seq analysis

Enhanced ChIP (eChIP) was performed as described (*58*), with minor modifications. Briefly, ∼0.1g of 2-week-old seedling was harvested, frozen in liquid nitrogen, and ground to a fine powder. Chromatin was crosslinked in 1.5 mL of fixation buffer (1xPBS, 1% (w/v) Formaldehyde, 1 mM Pefabloc SC, and 1xComplete Protease Inhibitor Cocktail) for 10 min at room temperature. Crosslink was quenched by adding 167 µL of 2M Glycine to a final concentration of 0.2 M and incubation for 5 min at room temperature. Samples were centrifuged at 3,000 x g at for 5 min at 4°C, and supernatant was discarded. Pellets were resuspended in 180 µL of Buffer S (50 mM HEPES-KOH (pH7.5), 150 mM NaCl, 1 mM EDTA, 1% Triton X-100, 0.1% sodium deoxycholate, 1% SDS, supplemented with cOmplete protease inhibitors) and incubated for 10 min at 4°C. Subsequently, 720 µL of Buffer F (50 mM HEPES-KOH (pH7.5), 150 mM NaCl, 1 mM EDTA, 1% Triton X-100, 0.1% sodium deoxycholate, supplemented with cOmplete protease inhibitors) was added. Chromatin was transferred to Covaris milliTUBE AFA Fiber and sonicated using Covaris system. Sonication was performed for 20 min (Duty Factor 5%, Peak Incident Power 140, and Cycles per Burst of 200) for histone modifications and for 15min (Duty Factor of 5%, Peak Incident Power of 105, and Cycles per Burst of 200) for 3xFLAG-tagged proteins. Sonicated samples were centrifuged at 20,000 x g for 10 min at 4°C, and supernatants were adjusted to a final volume of 1070 µL with Buffer F. Aliquot of chromatin were incubated overnight at 4°C with appropriate antibodies: anti-FLAG M2 (Sigma), anti-H3 (Abcam, ab1791), anti-H3K9me1 (Abcam, ab8896), and anti-H3K9me2 (MBL, mabi0317). Immunocomplexes were captured with 50 µL of Dynabeads Protein G (VERITAS), pre-washed twice with PBST (PBS with 0.1 % Tween 20), for ∼6 hr at 4°C with rotation. Beads were subsequently washed at 4°C for 10 min each with low-salt ChIP buffer (50 mM HEPES-KOH, 150 mM NaCl, 1 mM EDTA, 1% Triton X-100, 0.1% sodium deoxycholate, 0.1% SDS) three times, high-salt ChIP buffer (50 mM HEPES-KOH, 350 mM NaCl, 1 mM EDTA, 1% Triton X-100, 0.1% sodium deoxycholate, 0.1% SDS) twice, ChIP Wash Buffer (10 mM Tris-HCl pH 8.0, 250mM LiCl, 0.5% NP-40, 1 mM EDTA, 0.1% sodium deoxycholate) once, and TE Buffer once. Chromatin was eluted twice by incubation with 100 µL of ChIP elution buffer at 65°C for 15 min, followed by elution with 100 µL of 10 mM Tris-HCl (pH 8.5). Elutes were combined and reverse-crosslinked by overnight incubation with Proteinase K at 55°C. DNA was purified using the Monarch PCR & DNA Cleanup Kit (NEB) and used for library preparation with the ThruPLEX DNA-seq Kit (TaKaRa) and DNA Unique Dual Index Kit (Clontech).

### Data analysis

Sequenced reads were trimmed using Trimmomatic v0.33 (*59*). For methylome analysis, trimmed reads were mapped to Arabidopsis TAIR10 reference genome using Bismark (*60*), and generated CX reports were used for downstream analyses. DNA methylation levels across *Spm* transposons were visualized using deepTools (*61*). To identify differentially hypomethylated regions in CG context (CG hypoDMRs), DNA methylation levels were calculated in 50-bp bins and compared between TnpA transgenic plants and WT samples. Bins showing a decrease in CG methylation greater than 0.4 in TnpA transgenic plants relative to WT were defined as CG hypoDMRs. For TnpA^Spm3^, DNA sequences of CG hypoDMSs overlapping with *Spm3* were used for prediction of target motifs using DREME (*62*).

For eChIP-seq analysis, reads were mapped to Arabidopsis TAIR10 genome using Bowtie v1.1.2 (*63*). The resulting BAM files were used for peak calling with MACS3 v3.0.2, using two WT control samples for each transgenic sample (*64*). Shared peak regions of replicated ChIP-seq experiments were detected using the “insersect” command of BEDtools (*65*) and were used for estimation of target motifs with DREME for TnpA^Spm1^ and AtMu1c-TPase (*62*). Genomic localization of the predicted target motifs was identified using Bowtie (*63*). eChIP-seq read coverage was normalized to counts per million mapped reads (CPM) and aggregated over target transposons using deepTools (*61*). For ChIP-seq analysis of histone modifications, reads mapped to transposons were quantified using the “coverage” command of BEDtools with TAIR10 annotation as reference. RPKM+1 values for each histone modification were normalized by dividing by the corresponding H3 RPKM+1 values.

For RNA-seq analysis, trimmed reads were mapped to Arabidopsis TAIR10 reference genome using STAR (*66*). RPKM values for each gene were calculated from the ReadsPerGene.out.tab file generated using the “--quantMode GeneCounts” option. For fig. S3B, transposon-encoded genes showing a log2 fold change greater than 1.5 in transcript abundance [RPKM+1] in at least one of two RNA-seq replicates were counted as “active genes”.

### Phylogenetic analysis

For phylogenetic analysis, full-length transposon copies flanked by terminal inverted repeats (TIRs) and target site duplications (TSDs) were identified based on Arabidopsis TAIR10 genome annotation. For *Spm* transposons, full-length copies were selected according to the following criteria: (i) the presence of conserved CACTA sequences at both termini, (ii) a 3-bp TSD (allowing up to one nucleotide mismatch), (iii) the presence of transposon-encoded gene(s), and (iv) high expression in *ddm1* mutant (*42*). For a subset of *Spm* elements, genomic coordinates were manually corrected based on the precise positions of TIRs and TSDs. TIRs were detected and visualized using UGENE (*67*). For phylogenetic reconstruction, 120-bp sequences from 3’ terminal regions of *Spm* elements were used. For *AtMu* and *BRODYAGA2* transposons, copies longer than 1kb and flanked by TIRs and 8-11 bp TSDs (allowing up to one nucleotide mismatch) were selected. Although *AtMu1a* does not contain TSDs matching these criteria, its sequence was included because mobility of this copy has been reported previously (*43*) and was independently confirmed in this study (Fig. 4A and fig. S7A). Fragmented annotations in TAIR10 were manually merged when supported by TSD and TIR sequences. The orientations of merged *BRODYAGA2* copies were determined based on the longest annotated fragment and alignment with fixed sequences. For phylogenetic reconstruction, 120-bp sequences from 5’ terminal regions were used. Sequences were aligned using MUSCLE, and phylogenetic trees were constructed using the maximum likelihood neighbor-joining (NJ) method implemented in MEGA12 (*68*). Node support was assessed using 1,000 bootstrap replicates. List of transposons with corrected genomic coordinates are provided in Table S2 (*Spm*), S3 (*AtMu*), and S4 (*BRODYAGA2*), respectively.

### Detecting transposition mobilization

Identification of *de novo AtMu1* insertion from whole-genome resequencing data of self-pollinated *ddm1* mutants was performed as described in Fu *et al*. (*19*) (accession nos. DRA000420–000424). Briefly, reads containing *AtMu1*-specific terminal sequences were extracted using the following flanking sequences: GAGGGGGGCTCCTATAGGCA (5’ region of *AtMu1a*), AGGGGGCCTCCTATAGGCAG (3’ region of *AtMu1a*), GAGGGGGCCTCCTATAGGCA (5’ region of *AtMu1b*), AGGGGGCCTCCAATAGGTAG (3’ region of *AtMu1b*), GGTGACAGAGGCAGACAGCC (5’ region of *AtMu1c*), and GGGTGACAGAGGCAGGCAGC (3’ region of *AtMu1c*). The extracted flanking sequences were searched against the Arabidopsis TAIR10 genome using BLASTn, and the best unique hit with an E-value < 1e-5 was retained.

### Immunoblotting

Protein extraction and western blot analysis were performed as described previously (*69*). The anti-FLAG M2 antibody (Sigma-Aldrich) and an HRP-linked anti-mouse IgG antibody (Cytiva) were used as the primary and secondary antibodies at 1:5000 and 1:10000 dilutions, respectively. Signals were detected using ECL prime solution (Cytiva) with an iBright Imager.

### Identification of *Spm* DNA-binding domains

Protein sequences from *Spm* elements were retrieved from RepBase (https://www.girinst.org/repbase/). AlphaFold2 was used to model their 3D structure (*70*) and structural domains were defined for each protein as reported in (*40*). The DBDs of TnpA^Spm1^ and TnpA^Spm3^ were manually extracted based on previous data (*40*). Both DBDs, their complete structures and the TPase TnpD^Spm3^ were queried against all the RepBase *Spm* structural domains using foldseek (*39*), applying an e-value threshold of 0.1 and no prefiltering, and keeping only hits covering over 80% of the query length.

The predicted structural confidence (LDDT) for each Spm protein with a DBD hit fulfilling the above criteria was used to build a metaplot centered on the DBD hit. Similarly, splicing sites for each of these Spm proteins containing at least one splicing site were extracted from RepBase. These splicing sites were randomized 10,000 times maintaining the total number of sites per protein and used to build the random expectation.

To build the Uniform Manifold Approximation and Projection (UMAP) of all *Spm* structural domains, an exhaustive all-versus-all was performed. A distance metric defined as one minus the mean of TM-scores over the query and the target was applied in the UMAP projection.

In order to generate DBD phylogeny, a subset of proteins including a tri-helical Helix-turn-helix domain according to the Evolutionary Classification of Protein Domains Database (ECOD release 292.0) annotated as *Escherichia coli* insertion sequences were retrieved (*71*, *72*). These domain sequences, together with the Spm protein sequences spanning a hit on a DBD, were aligned using MAFFT E-INS-i (*73*), and a phylogenetic tree was built using iqtree3 using 1000 bootstrap replicates (*74*). All branches with support below 50% were collapsed into multifurcating nodes.

### Identification of *Spm* transcripts in long read sequencing data

Long read transcriptome data from zebrafish embryos at different development stages was downloaded from EBI. Protein sequences of zebrafish transposon proteins containing a DBD were used as queries in a tblastn search against all the long read transcriptomic raw reads. Raw reads were mapped to the Zebrafish Genome Reference Consortium GRCz11 (GCA_000002035) and overlapped with annotated transposable elements (TEs) with a length over 5kbp retrieved from a previous work (*41*). Blast hits overlapping annotated TEs were manually curated.

For Arabidopsis transposons, the TE location of the DBD was overlaid on a recently published TE annotation (*42*) and representative examples were selected.

### Identification of *MuDR* domestication events

All transposable elements with protein sequences from RepBase (https://www.girinst.org/repbase/) were retrieved and their 3D structure was modeled using AlphaFold2 implemented via ColabFold (*70*). High confidence proteins with experimental evidence at either transcript or protein level, excluding proteins annotated as TPases, were retrieved from UniProt and used to build a Foldseek database. A set of bacterial TPases were also retrieved to be used later as an outgroup.

To identify putative domesticated proteins, structural similarity with *AtMu1c* SWIM, WRKY and TPase domain was calculated using Foldseek search (*39*). RepBase proteins were analyzed in the same way. These domains were queried against the Foldseek database, and only hits fulfilling the following criteria were selected: 1) 80% of the query covered and with 90% probability to belong to the same protein fold as the query (qcov and prob fields reported by foldseek) or 2) 90% of the query covered and a TM-score over 0.5. The same parameters were applied to the RepBase proteins. Copy number was estimated for each putative domestication candidate by using its genomic coding sequence as a BLASTn query against the corresponding reference genome. Hits showing more than 90% nucleotide identity across more than 80% of the query length were counted. *MuDR*-related repetitive features were identified by running RepeatMasker on the 1-kb regions immediately upstream and downstream of each candidate locus. Candidates with no more than two additional high-identity matches and no *MuDR*-related sequences within 1 kb upstream or downstream of the candidate locus were classified as high-confidence domestication events.

Amino acid sequences of the TPase-like domains were aligned using MAFFT E-INS-i algorithm (*73*), and the phylogenetic tree was calculated using iqtree3 (*74*). Domestication events were identified by tree traversal and defined as clades of at least 90% of non-transposon proteins.

Hidden Markov Models for both domesticated and TPases were calculated from the alignment using Skylign (*75*). To calculate coordination distances for DDE-containing proteins, a triangle was defined positioning the vertices at the aspartate Cγ and the glutamate Cδ, both coinciding with the carbon of the carboxyl group on the side chain. Coordination distances were calculated as the maximum distance between the vertices and the triangle barycenter.

Visualization of 3D structural models was performed using ChimeraX (*76*).

### dN/dS estimation

To calculate the ratio of non-synonymous to synonymous mutations, the coding sequence for every analyzed protein was retrieved from RepBase and EBI for TPases and domesticated proteins, respectively. In the few cases where the coding sequences were not available, the corresponding protein was removed from the analyses. Protein sequences were aligned using MAFFT E-INS-i and the codon alignment was retrieved using PAL2NAL (*77*).

For each domestication event, the dN/dS ratio of individual proteins was estimated using the Goldman and Yang model (*78*) implemented in BioPython’s (*79*), comparing each protein with the closest TPase on the phylogenetic tree.

To estimate the per-branch dN/dS ratio on the phylogeny, a phylogenetic tree was built for each domestication event, including all the domesticated proteins and the ten closest TPases. The trees were built using iqtree3 as described above. Then, for each of these trees, PAML 4.10 program CODEML was used to test four different models (*80*). The first model set a single dN/dS fixed at 1 for the complete alignment; the second model estimated a single dN/dS for the complete alignment; the third model allowed different dN/dS for TPases and domesticated proteins, the formed being estimated and the latter being fixed at 1; and the fourth model allowed two different dN/dS, both of them estimated. Likelihood Ratio Tests were used to identify the best fit model for each domestication event.

## Acknowledgments

We thank members of the Quadrana group for discussions, Shusei Mori for technical advice on eChIP, Takumi Noyori and Soichi Inagaki for advice on data analysis, and Vincent Colot and Robert Martienssen for constructive comments on this manuscript. Computations were partially performed on the NIG supercomputer at ROIS National Institute of Genetics.

## Author contributions

Conceptualization: TS, LQ

Methodology: TS, CB, SF, ST, LQ

Investigation: TS, CB

Visualization: TS, CB, LQ

Funding acquisition: TS, LQ

Project administration: TS, LQ

Supervision: TS, LQ

Writing – original draft: TS, LQ

Writing – review & editing: TS, CB, SF, ST, LQ

## Data, code, and materials availability

New WGBS, ChIP-seq, and transcriptomic data has been deposited in the DNA Data Bank of Japan (DDBJ) under the accession number DRA026073. Publicly available WGBS and genome re-sequencing data reanalyzed in this study has been obtained from the ENA project accession PRJEB47214 and DDBJ project accession PRJDA60041, respectively.

## Supplementary Text supplementary text 1

Because DNA hypomethylation caused by *ddm1* mutation is heritable (*55*), *ddm1*-derived chromosomes can retain transcriptional activity after crossing with WT (*33*). Therefore, a potential transposon-encoded regulatory factor targeting *Spm1* should activate the Spm1-EGFP reporter in F1 progeny derived from crosses between *ddm1* and Spm1-EGFP plants. Consistent with this scenario, F1 progeny showed EGFP expression together with DNA hypomethylation (fig. S1B to D). The activated, hypomethylated state was heritable and persisted even in F2 plants homozygous for the WT *DDM1* allele, confirming the existence of a *trans*-acting regulatory system targeting *Spm1* (fig. S1C and D). Linkage mapping pinpointed a single locus responsible for *trans*-activation located in chromosome 2 (fig. S1E to G). This interval contains a full-length *Spm1* element, the same element used in the reporter construct.

## supplementary text 2

In plants, H3K9 methylation is a hallmark of constitutive heterochromatin (*81*). Interestingly, ChIP-seq analysis revealed that this modification persisted at target *Spm* elements even when they were transcriptionally activated by TnpA^Spm1^ (fig. S3A and B). These results indicate that TnpA-mediated anti-silencing mechanism can activate transposon-encoded genes without removing heterochromatic histone modifications. Since TnpA^Spm1^ primarily affects DNA methylation only at subterminal regions, we hypothesized that the active DNA demethylation pathway may be involved in TnpA^Spm1^-mediated anti-silencing. To test this hypothesis, we introduced TnpA^Spm1^ into triple mutants defective in the active DNA demethylation *ROS1*, *DML2* and *DML3* (*rdd*) carrying the Spm1-EGFP reporter (fig. S3C). While TnpA^Spm1^ activated Spm1-EGFP in WT plants, no EGFP signal was observed when transformed into *rdd* mutants (fig. S3C). Consistent with this, methylome analysis showed no change in DNA methylation in TnpA^Spm1^ *rdd* plants (fig. S3D). To exclude the possibility that the TnpA^Spm1^ transgene itself becomes silenced in *rdd* mutants (*82*), we also tested a transgene-free system leveraging *ddm1*-induced TnpA^Spm1^ activation (fig. S1B). F1 plants derived from a cross between *ddm1 rdd* and *rdd* carrying Spm1-EGFP reporter showed no EGFP signal, demonstrating that the active DNA demethylation pathway is essential for TnpA-mediated anti-silencing (fig. S3E). Notably, RT-PCR confirmed that TnpA^Spm1^ was still expressed in *ddm1 rdd* mutants and F1 plants (fig. S3F). Taken together, these results establish that TnpA^Spm1^-mediated anti-silencing requires the active DNA demethylation pathway.

The molecular mechanisms underlying active DNA demethylation differ substantially between plants (base-excision via DNA glycosylases) and animals (oxidative demethylation) (*82*). The dependency of TnpA^Spm1^ on plant-specific active DNA demethylases may reflect the evolution of anti-silencing factors adapted to plant epigenetic landscapes under relaxed evolutionary constraints.

## supplementary text 3

Although TnpA^Spm1^ binding was primarily detected at dense motif clusters within target *Spm* families, interestingly, we also identified a strong TnpA^Spm1^-binding locus outside *Spm* sequences in the pericentromeric region of chromosome 4 (fig. S4E). This locus comprises 22 tandemly repeated units of ∼2 kb sequences containing ∼5 motifs each, forming a 44 kb “motif cluster” we termed MC4 (motif cluster on chromosome 4) (fig. S4E). In TnpA^Spm1^ transgenic plants, MC4 is hypomethylated at CG sites, whereas non-CG methylation is increased. This locus is highly variable among natural Arabidopsis accessions, which harbor between 0 (C24) and 24 (Sha) copies of the tandem repeats (fig. S4F). Although its biological function is unknown, MC4 may have evolved via interaction with *Spm1* transposons.

## supplementary text 4

The Arabidopsis genome contains 12 *Spm* subfamilies. We hypothesized that additional families also encode family-specific anti-silencing factors. To test this, we analyzed WGBS data from sixteen *ddm1*-derived epigenetic recombinant inbred lines (epiRILs) to predict target motifs of *trans*-demethylation (*21*, *33*, *34*) (fig. S5A, see Materials and Methods). We identified 8-to 10-bp motifs for nine *Spm* families (fig. S5B). Hypomethylated regions over *Spm1* elements were enriched for the motif YCCTCGCAA (Y: C or T), closely matching the motif identified at TnpA^Spm1^ binding sites. Motif clustering for the nine *Spm* families revealed four groups, which mirrored the phylogeny of *Spm* consensus sequences, suggesting that diversification of anti-silencing specificity paralleled transposon diversification (fig. S5B). As observed for TnpA^Spm1^, single motifs are not sufficient for specificity, whereas clusters of four or more neighboring motifs are exclusively found on cognate or related copies, indicating that local accumulation of motifs provides an additional layer for anti-silencing specificity. In total, we identified at least four distinct anti-silencing systems among 12 *Spm* families (fig. S5B and C).

We next set out to test the regulatory activity of additional *Spm*-encoded systems. Among the 12 *Spm* families, *Spm3* harbors mobile copies in Arabidopsis (*28*-*30*), with AT2TE20205 representing the autonomous element (*28*). AT2TE20205 harbors a single gene that encodes both a predicted TnpD-like TPase and a TnpA-like factor through alternative splicing (*40*) (fig. S5D). We cloned the cDNA of the TnpA-like gene (designated TnpA^Spm3^) and transformed it into WT plants. Methylome analyses revealed TnpA^Spm3^-induced CG hypomethylation at *Spm3* elements, confirming its anti-silencing activity. Sequence motif enrichment analyses at TnpA^Spm3^-hypomethylated regions yielded “AYGCTAT”, which largely coincides with that predicted from the epiRIL analysis (fig. S5G and H). Altogether, our findings revealed the existence of multiple TnpA-mediated sequence-specific anti-silencing systems in the Arabidopsis genome.

## References and Notes

1. L. Sinzelle, Z. Izsvák, Z. Ivics. Molecular domestication of transposable elements: From detrimental parasites to useful host genes. Cell Mol Life Sci 66, 1073–1093 (2009). doi: 10.1007/s00018-009-8376-3; PMID: 19132291

2. R. L. Cosby, N. C. Chang, C. Feschotte. Host-transposon interactions: conflict, cooperation, and cooption. Genes Dev 33, 1098–1116 (2019). doi: 10.1101/gad.327312.119; PMID: 31481535

3. K. Naito, F. Zhang, T. Tsukiyama, H. Saito, C. N. Hancock, A. O. Richardson, Y. Okumoto, T. Tanisaka, S. R. Wessler. Unexpected consequences of a sudden and massive transposon amplification on rice gene expression. Nature 461, 1130–1134 (2009). doi: 10.1038/nature08479; PMID: 19847266

4. G. Kunarso, N. Y. Chia, J. Jeyakani, C. Hwang, X. Lu, Y. S. Chan, H. H. Ng, G. Bourque. Transposable elements have rewired the core regulatory network of human embryonic stem cells. Nat Genet 42, 631–634 (2010). doi: 10.1038/ng.600; PMID: 20526341

5. V. J. Lynch, R. D. Leclerk, G. May, G. P. Wagner. Transposon-mediated rewiring of gene regulatory networks contributed to the evolution of pregnancy in mammals. Nat Genet 43, 1154–1159 (2011). doi: 10.1038/ng.917; PMID: 21946353

6. A. Y. Du, J. D. Chobirko, X. Zhuo, C. Feschotte, T. Wang. Regulatory transposable elements in the encyclopedia of DNA elements. Nat Commun 15, 7594 (2024). doi: 10.1038/s41467-024-51921-6; PMID: 39217141

7. A. P. Marand, L. Jiang, F. Gomez-Cano, M. A. A. Minow, X. Zhang, J. P. Mendieta, Z. Luo, S. Bang, H. Yan, C. Meyer, L. Schlegel, F. Johannes, R. J. Schmitz. The genetic architecture of cell type-specific cis regulation in maize. Science 388, eads6601 (2025). doi: 10.1126/science.ads6601; PMID: 40245149

8. A. Balsalobre, J. Drouin. Pioneer factors as master regulators of the epigenome and cell fate. Nat Rev Mol Cell Biol 23, 449–464 (2022). doi: 10.1038/s41580-022-00464-z; PMID: 35264768

9. K. H. Burns. Our conflict with transposable elements and its implication for human disease. Annu Rev Pathol 15, 51–70 (2020). doi: 10.1146/annurev-pathmechdis-012419-032633; PMID: 31977294

10. M. E. Hudson, D. Lisch, P. H. Quail. The *FHY3* and *FAR1* genes encode transposon-related proteins involved in regulation of gene expression by the phytochrome A-signaling pathway. Plant J 34, 453–471 (2003). doi: 10.1046/j.1365-313x.2003.01741.x; PMID: 12753585

11. R. Lin, L. Ding, C. Casola, D. R. Ripoll, C. Feschotte, H. Wang. Transposase-derived transcription factors regulate light signaling in *Arabidopsis*. Science 318, 1302–1305 (2007). doi: 10.1126/science.1146281; PMID: 18033885

12. R. L. Cosby, J. Judd, R. Zhang, A. Zhong, N. Garry, E. J. Pritham, C. Feschotte. Recurrent evolution of vertebrate transcription factors by transposase capture. Science 371, eabc6405 (2021). doi: 10.1126/science.abc6405; PMID: 33602827

13. B. McClintock. The origin and behavior of mutable loci in maize. PNAS 36, 344–355 (1950). doi: 10.1093/genetics/38.6.579; PMID: 17247459

14. B. McClintock. Controlled mutation in maize. Carnegie Inst Wash Yr Bk 54, 245–255 (1955).

15. A. S. Klein, O. E. Nelson. Biochemical consequences of the insertion of a suppressor-mutator (*Spm*) receptor at the bronze-1 locus in maize. PNAS 80, 7591–7595 (1983). doi: 10.1073/pnas.80.24.7591; PMID: 16593396

16. O. E. Nelson, A. S. Klein. Characterization of an *Spm*-controlled bronze-mutable allele in maize. Genetics 106, 769–779 (1984). doi: 10.1093/genetics/106.4.769; PMID: 17246208

17. R. Martienssen, A. Barkan, W. C. Taylor, M. Freeling. Somatically heritable switches in the DNA modification of *Mu* transposable elements monitored with suppressible mutant in maize. Genes Dev 4, 331–343 (1990). doi: 10.1101/gad.4.3.331; PMID: 2159936

18. A. Barkan, R. A. Martienssen. Inactivation of maize transposon *Mu* suppresses a mutant phenotype by activating an outward-reading promoter near the end of *Mu1*. PNAS 88, 3502–3506 (1991). doi: 10.1073/pnas.88.8.3502; PMID: 1849660

19. Y. Fu, A. Kawabe, M. Etcheverry, T. Ito, A. Toyoda, A. Fujiyama, V. Colot, Y. Tarutani, T. Kakutani. Mobilization of a plant transposon by expression of the transposon-encoded anti-silencing factor. EMBO J 32, 2407–2417 (2013). doi: 10.1038/emboj.2013.169; PMID: 23900287

20. A. Hosaka, R. Saito, K. Takashima, T. Sasaki, Y. Fu, A. Kawabe, T. Ito, A. Toyoda, A. Fujiyama, Y. Tarutani, T. Kakutani. Evolution of sequence-specific anti-silencing systems in *Arabidopsis*. Nature Commun 8 2161 (2017). doi: 10.1038/s41467-017-02150-7; PMID: 29255196

21. T. Sasaki, K. Ro, E. Caillieux, R. Manabe, G. Bohl-Viallefond, P. Baduel, V. Colot, T. Kakutani, L. Quadrana. Fast co-evolution of anti-silencing systems and target sequences shapes the invasiveness of *Mu*-like DNA transposons. EMBO J 41, e110070 (2022). doi: 10.15252/embj.2021110070; PMID: 35285528

22. B. McClintock. Mutable loci in maize. Carnegie Inst Wash Yr Bk 50, 174–181 (1951).

23. P. Masson, G. Rutherford, J. A. Banks, N. Fedoroff. Essential large transcripts of the Maize *Spm* transposable element are generated by alternative splicing. Cell 58, 755–765 (1989). doi: 10.1016/0092-8674(89)90109-8; PMID: 2548734

24. M. Frey, J. Reinecke, S. Grant, H. Saedler, A. Gierl. Excision of the En/Spm transposable element of *Zea mays* requires two element-encoded proteins. EMBO J 9, 4037–4044 (1990). doi: 10.1002/j.1460-2075.1990.tb07625.x; PMID: 2174354

25. P. Masson, M. Strem, N. Fedoroff. The *tnpA* and *tnpD* gene products of the *Spm* element are required for transposition in tobacco. Plant Cell 3, 73–85 (1991). doi: 10.1105/tpc.3.1.73; PMID: 1668614

26. M. Schläppi, R. Raine, N, Fedoroff. Epigenetic regulation of the maize *Spm* transposable element: novel activation of a methylated promoter by TnpA. Cell 77, 427–437 (1994). doi: 10.1016/0092-8674(94)90157-0; PMID: 8181061

27. H. Cui, N. V. Fedoroff. Inducible DNA demethylation mediated by the maize Suppressor-mutator transposon-encoded TnpA protein. Plant Cell 14, 2883–2899 (2002). doi: 10.1105/tpc.006163; PMID: 12417708

28. A. Miura, S. Yonebayashi, K. Watanabe, T. Toyama, H. Shimada, T. Kakutani. Mobilization of transposons by a mutation abolishing full DNA methylation in Arabidopsis. Nature 411, 212–214 (2001). doi: 10.1038/35075612; PMID: 11346800

29. M. Catoni, T. Jonesman, E. Cerruti, J. Paszkowski. Mobilization of Pack-CACTA transposons in Arabidopsis suggests the mechanism of gene shuffling. Nucleic Acid Res 47, 1311–1320 (2019). doi: 10.1093/nar/gky1196; PMID: 30476196

30. L. Quadrana, M. Etcheverry, A. Gilly, E. Caillieux, M. A. Madoui, J. Guy, A. Bortolini Silveira, S. Engelen, V. Baillet, P. Wincker, J. M. Aury, V. Colot. Transposition favors the generation of large effect mutations that may facilitate rapid adaptation. Nature Commun 10, 3421 (2019). doi: 10.1038/s41467-019-11385-5; PMID: 31366887

31. K. M. Creasey, J. Zhai, F. Borges, F. Van Ex, M. Regulski, B. C. Meyers, R. A. Martienssen. miRNAs trigger widespread epigenetically activated siRNAs from transposons in *Arabidopsis*. Nature 508, 411–415 (2014). doi: 10.1038/nature13069; PMID: 24670663

32. T. Sasaki, K. Kato, A. Hosaka, Y. Fu, A. Toyoda, A. Fujiyama, Y. Tarutani, T. Kakutani. Arms race between anti-silencing and RdDM in noncoding regions of transposable elements. EMBO Rep 24, e56678 (2023). doi: 10.15252/embr.202256678; PMID: 37272687

33. F. Johannes, E. Porcher, F. K. Teixeira, V. Saliba-Colombani, M. Simon, A. Bulski, J. Albuisson, F. Heredia, P. Audigier, D. Bouchez, C. Dillmann, P. Guerche, F. Hospital, V. Colot. Assessing the impact of transgenerational epigenetic variation on complex traits. PLoS Genet 5, e1000530 (2009). doi: 10.1371/journal.pgen.1000530; PMID: 19557164

34. P. Baduel, L. D. Olivier, E. Cortex, G. Bohl-Viallefond, C. Xu, M. El Messaoudi, A. Petit, M. Draï, M. Barois, V. Singh, A. Sarazin, F. K. Texeira, M. Boccare, E. Gilbault, A. de France, L. Quadrana, O. Loudet, V. Colot. Transposable elements are vectors of recurrent transgenerational epigenetic inheritance. Science 390, eady3475 (2025). doi: 10.1126/science.ady3475; PMID: 40966307

35. H. Saze, T. Kakutani. Heritable epigenetic mutation of a transposon-flanked *Arabidopsis* gene due to lack of the chromatin-remodeling factor DDM1. EMBO J 26, 3641–3652 (2007). doi: 10.1038/sj.emboj.7601788; PMID: 17627280

36. I. R. Henderson, S. E. Jacobsen. Tandem repeats upstream of the *Arabidopsis* endogene SDC recruit non-CG methylation and initiate siRNA spreading. Genes Dev 22, 1597–1606 (2008). doi: 10.1101/gad.1667808; PMID: 18559476

37. T. Sasaki, A. Kobayashi, H. Saze, T. Kakutani. RNAi-independent *de novo* DNA methylation revealed in Arabidopsis mutants of chromatin remodeling gene *DDM1*. Plant J 70, 750–758 (2012). doi: 10.1111/j.1365-313X.2012.04911.x; PMID: 22269081

38. M. Ong-Abdullah, J. M. Ordway, N. Jiang, S. E. Ooi, S. Y. Kok, N. Sarpan, N. Azimi, A. T. Hashim, Z. Ishak, S. K. Rosli, F. A. Malike, N. A. Bakar, M. Marjuni, N. Abdullah, Z. Yaakub, M. D. Amiruddin, R. Nookiah, R. Singh, E. T. Low, K. L. Chan, N. Azizi, S. W. Smith, B. Bacher, M. A. Budiman, A. Van Brunt, C. Wischmeyer, M. Beil, M. Hogan, N. Lakey, C. C. Lim, X. Arulandoo, C. K. Wong, C. N. Choo, W. C. Wong, Y. Y. Kwan, S. S. Alwee, R. Sambanthamurthi, R. A. Martienssen. Loss of Karma transposon methylation underlies the mantled somaclonal variant of oil palm. Nature 525, 533–537 (2015). doi: 10.1038/nature15365; PMID: 26352475

39. M. van Kempen, S. S. Kim, C. Tumescheit, M. Mirdita, J. Lee, C. L. M. Gilchrist, J. Söding, M. Steinegger. Fast and accurate protein structure search with Foldseek. Nat Biotechnol 42, 243–246 (2024). doi: 10.1038/s41587-023-01773-0; PMID: 37156916

40. C. Borredá, P. Vendrell-Mir, B. Leduque, M. Bueno Merino, C. Oury, N, Glab, V. Colot, L. Quadrana. A functional atlas of transposon-encoded products and their integration into host networks. Nat Commun 10.1038/s41467-026-76280-2 (2026).

41. N.-C. Chang, Q. Rovira, J. Walls, C. Feschotte, J. M. Vaquerizas. Zebrafish transposable elements show extensive diversification in age, genomic distribution, and developmental expression. Genome Res 32, 1408–1423 (2022). doi: 10.1101/gr.275655.121; PMID: 34987056

42. S. Oberlin, A. Sarazin, C. Chevalier, O. Voinnet, A. Marí-Ordóñez. A genome-wide transcriptome and translatome analysis of *Arabidopsis* transposons identifies a unique and conserved genome expression strategy for *Ty1*/*Copia* retroelements. Genome Res 27, 1549–1562 (2017). doi: 10.1101/gr.220723.117; PMID: 28784835

43. T. Singer, C. Yordan, R. A. Martienssen. Robertson’s *Mutator* transposons in *A. thaliana* are regulated by the chromatin-remodeling gene *Decrease in DNA Methylation* (*DDM1*). Genes Dev 15, 591–602 (2001). doi: 10.1101/gad.193701; PMID: 11238379

44. S. Tsukahara, A. Kobayashi, A. Kawabe, O. Mathieu, A. Miura, T. Kakutani. Bursts of retrotransposition reproduced in Arabidopsis. Nature 461, 423–426 (2009). doi: 10.1038/nature08351; PMID: 19734880

45. T. Kabelitz, C. Kappel, K. Henneberger, E. Benke, C. Nöh, I. Bäurle. eQTL mapping of transposon silencing reveals a position-dependent stable escape from epigenetic silencing and transposition of *AtMu1* in the *Arabidopsis* lineage. Plant Cell 26, 3261–3271 (2014). doi: 10.1105/tpc.114.128512; PMID: 25096782

46. H. Quesneville. Twenty years of transposable element analysis in the *Arabidopsis thaliana* genome. Mob DNA 11, 28 (2020). doi: 10.1186/s13100-020-00223-x

47. Z. Joly-Lopez, E. Forczek, D. R. Hoen, N, Juretic, T. E. Bureau. A gene family derived from transposable elements during early angiosperm evolution has reproductive fitness benefits in *Arabidopsis thaliana*. PLoS Genet 8, e1002931 (2012). doi: 10.1371/journal.pgen.1002931; PMID: 22969437

48. Z. Joly-Lopez, D. R. Hoen, M. Blanchette, T. E. Bureau. Phylogenetic and genomic analyses resolve the origin of important plant genes derived from transposable elements. Mol Biol Evol 33, 1937–1956 (2016). doi: 10.1093/molbev/msw067; PMID: 27189548

49. D. Lisch, P. Chomet, M. Freeling. Genetic characterization of the *mutator* system in maize: behavior and regulation of *mu* transposons in minimal line. Genetics 139, 1777–1796 (1995). doi: 10.1093/genetics/139.4.1777; PMID: 7789777

50. D. Burgess, H. Li, M. Zhao, S. Y. Kim, D. Lisch. Silencing of *mutator* elements in maize involves distinct populations of small RNAs and distinct patterns of DNA methylation. Genetics 215, 379–391 (2020). doi: 10.1534/genetics.120.303033; PMID: 32229532

51. D. Schwartz. Gene-controlled cytosine demethylation in the promoter region of the *Ac* transposable element in maize. Proc Natl Acad Sci USA 86, 2789–2793 (1989). doi: 10.1073/pnas.86.8.2789; PMID: 16594028

52. S. N. Hashida, K. Kitamura, T. Mikami, Y. Kishima. Temperature shift coordinately changes the activity and the methylation state of transposon Tam3 in *Antirrhium majus*. Plant Physiol 132, 1207–1216 (2003). doi: 10.1104/pp.102.017533; PMID: 12857803

53. T. Kawashima, F. Berger. Epigenetic reprogramming in plant sexual reproduction. Nat Rev Genet 15, 613–624 (2014). doi: 10.1038/nrg3685; PMID: 25048170

54. R. K. Cowan, D. R. Hoen, D. J. Schoen, T. E. Bureau. MUSTANG is a novel family of domesticated transposase genes found in diverse angiosperms. Mol Biol Evol 22, 2084–2089 (2005). doi: 10.1093/molbev/msi202; PMID: 15987878

55. A. Vongs, T. Kakutani, R. A. Martienssen, E. J. Richards. Arabidopsis thaliana DNA methylation mutants. Science 260, 1926–1928 (1993). doi: 10.1126/science.8316832; PMID: 8316832

56. J. M. Alonso, A. N. Stepanova, T. J. Leisse, C. J. Kim, H. Chen, P. Shinn, D. K. Stevenson, J. Zimmerman, P. Barajas, R. Cheuk, C. Gadrinab, C. Heller, A. Jeske, E. Koesema, C. C. Meyers, H. Parker, L. Prednis, Y. Ansari, N. Choy, H. Deen, M. Geralt, N. Hazari, E. Hom, M. Karnes, C. Mulholland, R. Ndubaku, I. Schmidt, P. Guzman, L. Aguilar-Henonin, M. Schmid, D. Weigel, D. E. Carter, T. Marchand, E. Risseeuw, D. Brogden, A. Zeko, W. L. Crosby, C. C. Berry, J. R. Ecker. Genome-wide insertional mutagenesis of *Arabidopsis thaliana*. Science 301, 653–657 (2003). doi: 10.1126/science.1086391; PMID: 12893945

57. S. J. Clough, A. F. Bent. Floral dip: a simplified method for *Agrobacterium*-mediated transformation of *Arabidopsis thaliana*. Plant J 16, 735–743 (1998). doi: 10.1046/j.1365-313x.1998.00343.x; PMID: 10069079

58. L. Zhao, L. Xie, Q. Zhang, W. Ouyan, L. Deng, P. Guan, M. Ma, Y. Li, Y. Zhang, Q. Xioa, J. Zhang, H. Li, S. Wang, J. Man, Z. Cao, Q. Zhang, Q. Zhang, G. Li, X. Li. Integrative analysis of reference epigenomes in 20 rice varieties. Nat Commun 11, 2658 (2020). doi: 10.1038/s41467-020-16457-5; PMID: 32461553

59. A. M. Bolger, M. Lohse, B. Usadel. Trimmomatic: a flexible trimmer for Illumina sequence data. Bioinformatics 30, 2114–2120 (2014). doi: 10.1093/bioinformatics/btu170; PMID: 24695404

60. F. Krueger F, S. R. Andrews. Bismark: a flexible aligner and methylation caller for Bisulfite-seq applications. Bioinformatics 27, 1571–1572 (2011). doi: 10.1093/bioinformatics/btr167; PMID: 21493656

61. F. Ramírez, D. P. Ryan, B. Grüning, V. Bhardwaj, F. Kilpert, A. S. Richter, S. Heyne, F. Dündar, T. Manke. deepTools2: a next generation web server for deep-sequencing data analysis. Nucleic Acids Res 44, W160–165 (2016). doi: 10.1093/nar/gkw257; PMID: 27079975

62. T. L. Bailey. DREME: motif discovery in transcription factor ChIP-seq data. Bioinformatics 27, 1653–1659 (2011). doi: 10.1093/bioinformatics/btr261; PMID: 21543442

63. B. Langmead, C. Trapnell, M. Pop, S. L. Salzberg. Ultrafast and memory-efficient alignment of short DNA sequences to the human genome. Genome Biology 10, R25 (2009). doi: 10.1186/gb-2009-10-3-r25; PMID: 19261174

64. Y. Zhang, T. Liu, C. A. Meyer, J. Eeckhoute, D. S. Johnson, B. E. Bernstein, C. Nusbaum, R. M. Myers, M. Brown, W. Li, X. S. Liu. Model-based analysis of ChIP-seq (MACS). Genome Biol 9, R137 (2008). doi: 10.1186/gb-2008-9-9-r137; PMID: 18798982

65. A. R. Quinlan, I. M. Hall. BEDTools: a flexible suite of utilities for comparing genomic features. Bioinformatics 26, 841–842 (2010). doi: 10.1093/bioinformatics/btq033; PMID: 20110278

66. A. Dobin, C. A. Davis, F. Schlesinger, J. Drenkow, C. Zaleski, S. Jha, P. Batut, M. Chaisson, T. R. Gingeras. STAR: ultrafast universal RNA-seq aligner. Bioinformatics 29, 15–21 (2013). doi: 10.1093/bioinformatics/bts635; PMID: 23104886

67. K. Okonechnikov, O. Golosova, M. Fursov; UGENE team. Unipro UGENE: a unified bioinformatics toolkit. Bioinformatics 28, 1166–1167 (2012). doi: 10.1093/bioinformatics/bts091; PMID: 22368248

68. S. Kumar, G. Stecher, M. Suleski, M. Sanderford, S. Sharma, K. Tamura. MEGA12: Molecular evolutionary genetic analysis version 12 for adaptive and green computing. Mol Biol Evol 41, msae263 (2024). doi: 10.1093/molbev/msae263; PMID: 39708372

69. T. Sasaki, U. Naumann, P. Forai, A. J. Matzke, M. Matzke. Unusual case of apparent hypermutation in *Arabidopsis thaliana*. Genetics 192, 1271–1280 (2012). doi: 10.1534/genetics.112.144634; PMID: 23023006

70. M. Mirdita, K. Schütze, Y. Moriwaki, L. Heo, S. Ovchinnikov, M. Steinegger. ColabFold: making protein folding accessible to all. Nat Methods 19, 679–682 (2022). doi: 10.1038/s41592-022-01488-1; PMID: 35637307

71. H. Cheng, R. D. Schaeffer, Y. Liao, L. N. Kinch, J. Pei, S. Shi, B. H. Kim, N. V. Grishin. ECOD: an evolutionary classification of protein domains. PLoS Comput Biol 10, e1003926 (2014). doi: 10.1371/journal.pcbi.1003926; PMID: 25474468

72. R. D. Schaeffer, K. D. Medvedev, A. Andreeva, S. R. Chuguransky, B. L. Pinto, J. Zhang, Q. Cong, A. Bateman, N. V. Grishin. ECOD: integrating classification of protein domains from experimental and predicted structures. Nucleic Acid Res 53: D411–D418 (2025). doi: 10.1093/nar/gkae1029; PMID: 39565196

73. K. Katoh, D. M. Standley. MAFFT multiple sequence alignment software version 7: improvements in performance and usability. Mol Biol Evol 30, 772–780 (2013). doi: 10.1093/molbev/mst010; PMID: 23329690

74. T. K. F. Wong, N. Ly-Trong, H. Ren, P. Demotte, H. Baños, A. J. Roger, E. Susko, C. Bielow, N. De Maio, N. Goldman, M. W. Hahn, M. dos Reis, L. S. Vinh, G. Huttley, R. Lanfear, B. Q. Minh. IQ-TREE 3: phylogenomic interface software using complex evolutionary models. Mol Biol Evol 43, msag117 (2026). doi: 10.1093/molbev/msag117; PMID: 42085559

75. T. J. Wheeler, J. Clements, R. D. Finn. Skylign: a tool for creating informative, interactive logos representing sequence alignments and profile hidden Markov models. BMC Bioinformatics 15, 7 (2014). doi: 10.1186/1471-2105-15-7; PMID: 24410852

76. E. C. Meng, T. D. Goddard, E. F. Petterson, G. S. Couch, Z. J. Pearson, J. H. Morris, T. E. Ferrin. UCSF ChimeraX: Tools for structure building and analysis. Protein Sci 32, e4792 (2023). doi: 10.1002/pro.4792; PMID: 37774136

77. M. Suyama, D. Torrents, P. Bork. PAL2NAL: robust conversion of protein sequence alignments into the corresponding codon alignments. Nucleic Acid Res 34, W609–W612 (2006). doi: 10.1093/nar/gkl315; PMID: 16845082

78. N. Goldman, Z. Yang. A codon-based model of nucleotide substitution for protein-coding DNA sequences. Mol Biol Evol 11, 725–736 (1996). doi: 10.1093/oxfordjournals.molbev.a040153; PMID: 7968486

79. P. J. Cock, T. Antao, J. T. Chang, B. A. Chapman, C. J. Cox, A. Dalke, I. Friedberg, T. Hamelryck, F. Kauff, B. Wilczynski, M. J. de Hoon. Biopython: freely available Python tools for computational molecular biology and bioinformatics. Bioinformatics 25, 1422–1423 (2009). doi: 10.1093/bioinformatics/btp163; PMID: 19304878

80. Z. Yang. PAML 4: phylogenetic analysis by maximum likelihood. Mol Biol Evol 24, 1586–1591 (2007). doi: 10.1093/molbev/msm088; PMID: 17483113

81. A. V. Gendrel, Z. Lippman, C. Yordan, V. Colot, R. A. Martienssen. Dependence of heterochromatic histone H3 methylation patterns on the Arabidopsis gene DDM1. Science 297, 1871–1873 (2002). doi: 10.1126/science.1074950; PMID: 12077425

82. J. K. Zhu. Active DNA demethylation mediated by DNA glycosylases. Annu Rev Genet 43, 143–166 (2009). doi: 10.1146/annurev-genet-102108-134205; PMID: 19659441

83. W. Tian, R. Wang, C. Bo, Y. Yu, Y. Zhang, G.-I. Shin, W.-Y. Kim, L. Wang. *SDC* mediates DNA methylation-controlled clock pace by interacting with ZTL in *Arabidopsis*. Nucleic Acid Res 49, 3764–3780 (2021). doi: 10.1093/nar/gkab128; PMID: 33675668

